# Cave *Thiovulaceae* differ metabolically and genomically from marine species

**DOI:** 10.1101/2020.11.04.367730

**Authors:** Mina Bizic, Traian Brad, Danny Ionescu, Lucian Barbu-Tudoran, Joost Aerts, Radu Popa, Luca Zoccarato, Jessica Ody, Jean-François Flot, Scott Tighe, Daniel Vellone, Serban M. Sarbu

**Affiliations:** Leibniz Institute for Freshwater Ecology and Inland Fisheries, IGB, Dep 3, Experimental Limnology, Zur Alte Fischerhütte 2, OT Neuglobsow, 16775 Stechlin, Germany; Berlin-Brandenburg Institute of Advanced Biodiversity Research (BBIB), Berlin, Germany; “Emil Racoviţă” Institute of Speleology, Clinicilor 5-7, 400006 Cluj-Napoca Romania; Institutul Român de Ştiinţă şi Tehnologie, Virgil Fulicea nr. 3, 400022 Cluj-Napoca, Romania; Center for Electron Microscopy, “Babeş-Bolyai” University, Clinicilor 5, 400006 Cluj-Napoca, Romania; Department of Molecular Cell Physiology, Faculty of Earth and Life sciences, De Boelelaan 1085, 1081 HV Amsterdam, The Netherlands; River Road Research, 62 Leslie St. Buffalo, NY 1421, USA; Evolutionary Biology and Ecology, Université libre de Bruxelles (ULB), C.P. 160/12, Avenue F.D. Roosevelt 50, 1050 Brussels, Belgium; Interuniversity Institute of Bioinformatics in Brussels – (IB)^2^, Brussels, Belgium; Vermont Integrative Genomics Lab, University of Vermont Cancer Center, Health Science Research Facility, Burlington, Vermont, 05405, USA; “Emil Racoviţă” Institute of Speleology, Frumoasa 31-B, 010986 Bucureşti, Romania; Department of Biological Sciences, California State University, Chico 95929, USA

**Keywords:** *Thiovulum*, sulfur, DNRA, Movile Cave, sulfide-oxidation

## Abstract

Life in Movile Cave (Romania) relies entirely on carbon fixation by bacteria. The microbial community in the surface water of Movile Cave’s hypoxic air bells is dominated by large spherical-ovoid bacteria we identified as *Thiovulum* sp. (*Campylobacterota*). These form a separate phylogenetic cluster within the *Thiovulaceae*, consisting mostly of freshwater cave bacteria. We compared the closed genome of this *Thiovulum* to that of the marine strain *Thiovulum* ES, and to a genome we assembled from public data from the sulfidic Frasassi caves. The Movile and Frasassi *Thiovulum* were very similar, differing greatly from the marine strain. Based on their genomes, cave *Thiovulum* can switch between aerobic and anaerobic sulfide oxidation using O_2_ and NO_3_^-^ as electron acceptors, respectively. NO_3_^-,^ is likely reduced to NH_3_ via dissimilatory nitrate reduction to ammonia using periplasmic nitrate reductase (Nap) and hydroxylamine oxidoreductase. Thus, *Thiovulum*, is likely important to both S and N cycles in sulfidic subterranean aquatic ecosystems. Additionally, we suggest that the short peritrichous flagella-like structures typical of *Thiovulum* are type IV pili, for which genes were found in all *Thiovulum* genomes. These pili may play a role in veil formation, connecting adjacent cells and the exceptionally fast swimming of these bacteria.

## INTRODUCTION

Movile Cave is located near the town of Mangalia, SE Romania (43°49’32”N, 28°33’38”E), 2.2 km inland from the Black Sea shore. It consists of a 200 m long upper dry passage that ends in a small lake allowing access to a 40 m long, partially submerged lower cave level (Fig. 1). Thick and impermeable layers of clays and loess cover the limestone in which the cave is developed, preventing input of water and nutrients from the surface (Lascu *et al*., 1994). Sulfidic groundwater flows constantly at the bottom of Movile Cave’s lower passages. Because of the morphology of the lower cave passages (Fig. 1) and a slight difference in water temperatures, the water near the surface is practically stagnant. Oxygen penetrates up to 1 mm of the water column, below which the water is anoxic (Riess *et al*., 1999).

**Figure 1.**
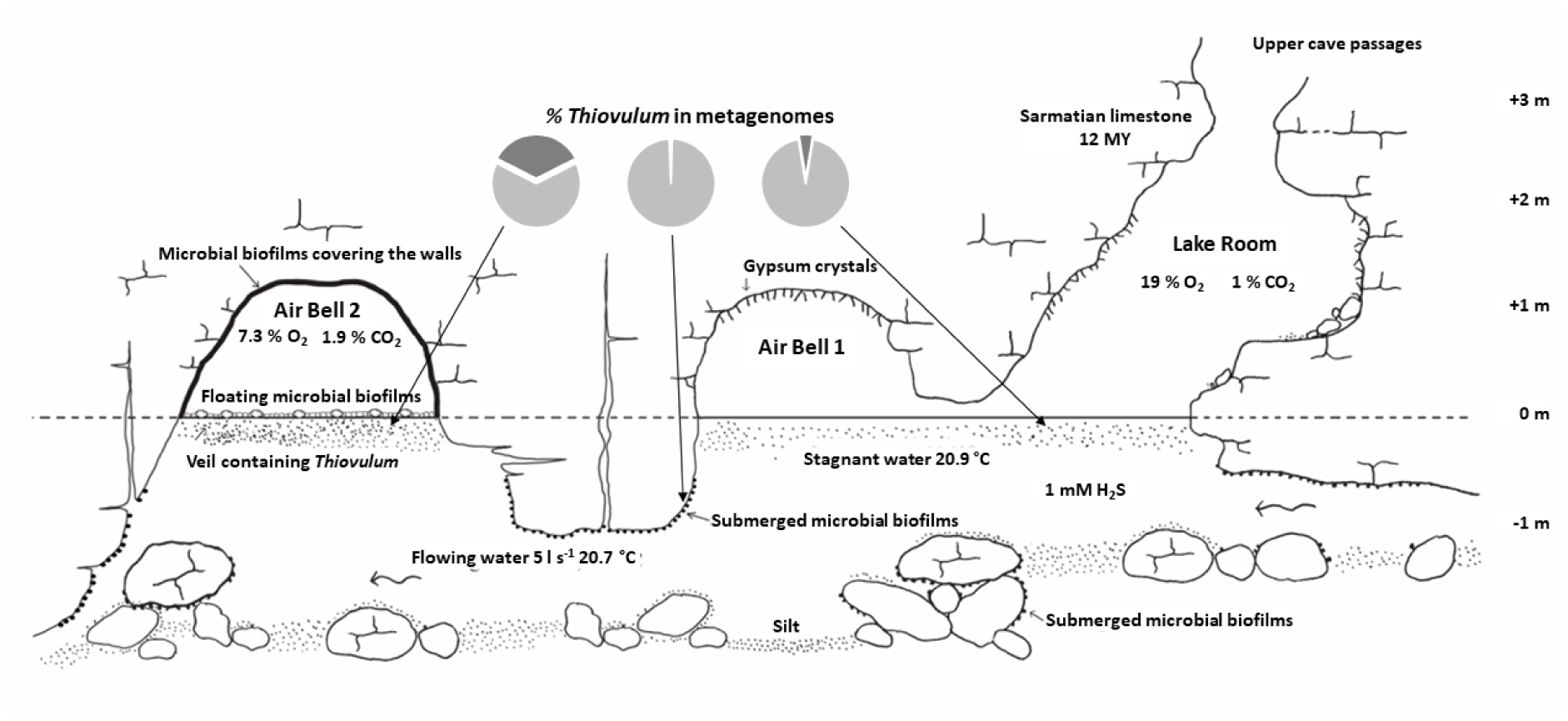
Longitudinal profile of the sampling area in Movile Cave (modified after Sarbu and Popa, 1992). The microbial community containing *Thiovulum* cells (depicted here as dots present beneath the water surface) was sampled in the Lake Room and in Air Bells 1 and 2. *Thiovulum* 16S rRNA made up 5 %, 0.9 % and 35 % of the 16S rRNA genes retrieved from metagenomic samples (dark gray in pies) from these cave sections, respectively. More details on community composition are presented in Supplementary Figure 1.

Cave ecosystems, normally characterized by stable conditions, provide a window into subsurface microbiology (Engel, 2015). In the absence of natural light, these ecosystems are typically fueled by chemolithoautotrophy via the oxidation of reduced compounds such as H_2_S, Fe^2+^, Mn^2+^, NH_3_, CH_4_, and H^+^. Most of the microbiological studies performed in Movile Cave (summarized in (Kumaresan *et al*., 2014) are based on samples of microbial biofilms floating on the water surface or covering rock surfaces in the cave’s Air Bells (Fig. 1), where the atmosphere is low in O_2_ (7-10 %) and enriched in CO_2_ (2.5 %) and CH_4_

(1-2 %) (Sarbu, 2000). There, chemoautotrophic microorganisms, living at the water surface, oxidize reduced chemical compounds such as H_2_S, CH_4_ and NH_4_^+^ from the thermo-mineral groundwater (Sarbu, 2000; Sarbu and Kane, 1995; Sarbu *et al*., 1996). *Thiobacillus, Thiothrix, Thioploca, Thiomonas* and *Sulfurospirillum* oxidize H_2_S using O_2_ or NO_3_^-^ as electron acceptors (Rohwerder *et al*., 2003; Chen *et al*., 2009; Flot *et al*., 2014). The methanotrophs *Methylomonas, Methylococcus* and *Methylocystis* (Hutchens *et al*., 2004), *Methanobacterium* (Schirmack *et al*., 2014) and *Methanosarcina* (Ganzert *et al*., 2014) are also found in the cave, alongside other methylothrophs such as *Methylotenera, Methylophilus* and *Methylovorus* (Rohwerder *et al*., 2003; Chen *et al*., 2009). Chen *et al*. (2009) further identified in this cave ammonia and nitrite oxidizers from the genera *Nitrospira* and *Nitrotoga*.

In the lower level of Movile Cave, directly below the water surface (not deeper than 2-3 mm) we observed a loose floating veil resembling a slow-moving white cloud (Fig. 2 and Supplementary video 1). Using genetic and microscopic analysis, we concluded that this underwater agglomeration of bacteria is dominated by a species of the genus *Thiovulum* (Fig. 1, S1 and results).

**Figure 2.**
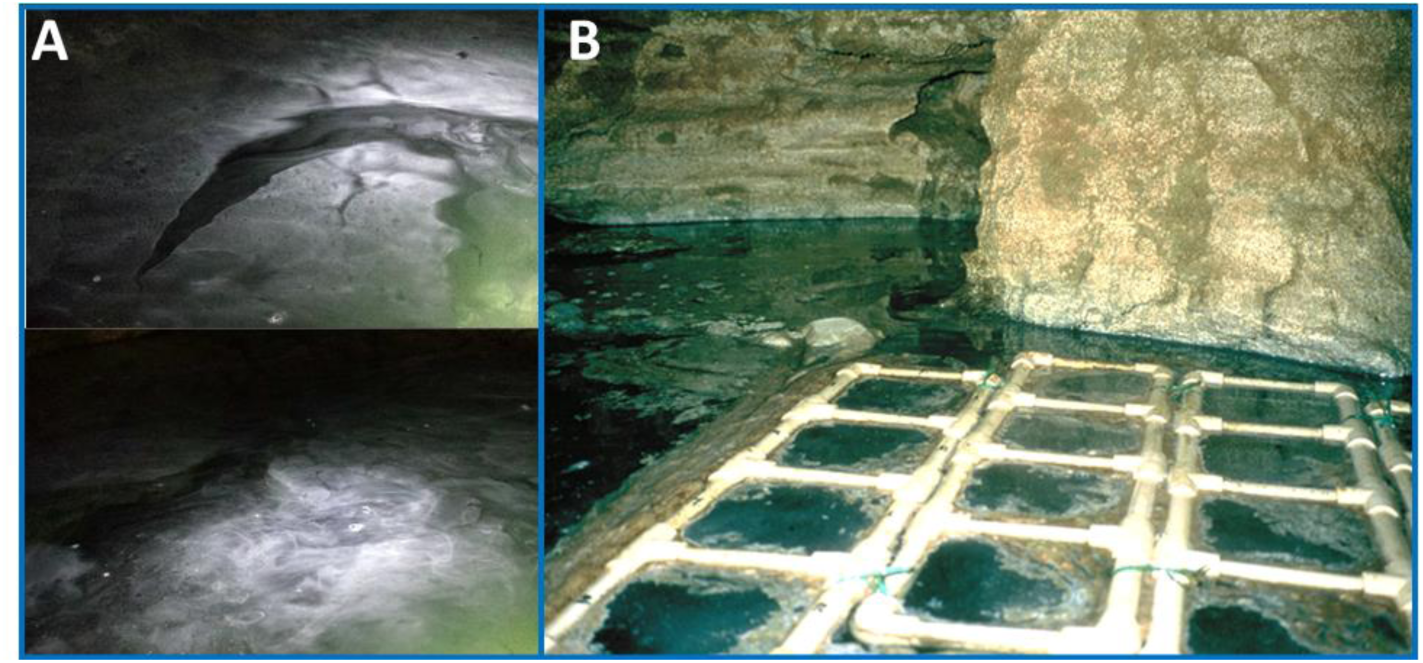
Images of subsurface veil from the Lake Room (A) and Air Bell 2 (B) in Movile Cave. See also Supplementary Video 1.

*Thiovulum* is a large bacterium, typically < 25 μm in diameter (Robertson *et al*., 2015) but can reach 45 μm (Sylvestre *et al*., 2021). It is a sulfur-oxidizing chemolithoautotrophic bacteria (Wirsen and Jannasch, 1978) with an extremely fast motility (Garcia-Pichel, 1989; Thar and Fenchel, 2001). *Thiovulum* is known to form a veil close to surfaces (Petroff *et al*., 2015; Robertson *et al*., 2015), to which it can attach through a secreted stalk (De Boer *et al*., 1961). It is normally located close to the oxic-anoxic interface near sediments or microbial mats (Marshall *et al*., 2012; Robertson *et al*., 2015; Jorgensen and Revsbech, 1983) where the 2D organization of the veil and the rapid movements of the cells’ flagella produce a convective transport of O_2_ (Fenchel and Glud, 1998).

To the best of our knowledge, this is the first description of fully planktonic *Thiovulum* swarms/veils at distance from any solid surface. Here we provide further morphological and genomic information on this bacterium, offering new insights into its metabolic properties and raising novel questions.

## MATERIALS AND METHODS

A detailed description of the methods is provided in the supplementary material.

Replicate samples of water were collected into sterile containers from the surface of the small sulfidic lake and from the Air Bells in the lower section of Movile Cave (Fig. 1). Unpreserved 50 ml water samples were immediately brought to the laboratory and inspected using optical microscopy. Samples for DNA/RNA analysis were preserved in ethanol (final concentration of 50 %). Additionally, samples were preserved with formaldehyde (final concentration of 4 %) for cell enumeration. Samples for electron microscopy were fixed with 2.7 % glutaraldehyde in phosphate buffered saline (1× PBS). Samples for DNA extraction were collected in July 2019 whereas samples for RNA extraction were collected in August 2021.

### Electron microscopy and elemental analysis

Fixed, dehydrated, and epoxy-embedded samples were sliced (100 nm thickness), stained with lead citrate and uranyl acetate (Hayat, 2001 and analyzed with Jeol JEM transmission electron microscope (Jeol, Japan). Samples for scanning electron microscopy (SEM) were sputtered with gold and examined on a JEOL JSM 5510 LV microscope (Jeol, Japan). Energy-dispersive X-ray spectroscopy (EDX) analysis was performed with an EDX analyzer (Oxford Instruments, Abingdon, UK) and with the INCA 300 software.

### DNA and RNA Extraction

Genomic DNA was extracted using a modified version of the Omega BioTek Universal Metagenomics kit protocol (OMEGA Bio-Tek, GA, USA) (see supplementary material) from samples filtered on with a 0.2 μm Isopore membrane filter (Millipore Sigma, MA, USA).

For RNA extraction, total nucleic acids were extracted from polycarbonate filters (Millipore, 0.2 μm pore size) following Nercessian et *al*. (2005) with minor modifications (see supplementary material. DNA was digested by two sequential treatments with the TurboDNAfree Kit (Invitrogen ThermoFisher Scientific, Dreieich, Germany) following the manufacturer’s instructions. DNA removal was evaluated using a PCR reaction for 16S rRNA gene. First strand cDNA was then generated using the High-Capacity cDNA Reverse Transcription Kit (Applied Biosciences, ThermoFisher scientific), and was sent for sequencing at the Core Genomic Facility at RUSH university, Chicago, IL, USA.

### 16S rRNA gene amplicon sequencing and processing

PCR reactions were performed in triplicates targeting the V3-V4 region of the 16S rRNA gene, using the V3 forward primer S-D-Bact-0341-b-S-17, 5’-CCTACGGGNGGCWGCAG-3 (Herlemann *et al*., 2011), and the V4 reverse primer S-D-Bact-0785-a-A-21, 5’-GACTACHVGGGTATCTAATCC-3 (Muyzer *et al*., 1993), resulting in fragments of ∼430 bp. The primers were dual barcoded in a way compatible with Illumina sequencing platforms (as described in Caporaso *et al*. (2011).

Composite samples were paired-end sequenced at the Vrije Universiteit Amsterdam Medical Center (Amsterdam, The Netherlands) on an Illumina MiSeq Sequencer. The paired sequences were dereplicated using the dedupe tool of the BBTools package (sourceforge.net/projects/bbmap/) aligned and annotated using the SINA aligner (Pruesse *et al*., 2012) against the SILVA SSU database (v 138.1) (Quast *et al*., 2013)

A maximum-likelihood phylogenetic tree was calculated including only long 16S rRNA sequences, using FastTree 2 (Price *et al*., 2010) using all *Thiovulum* sequences in the SILVA database (n=71), three 16S rRNA sequences obtained from the assembled genome of the Movile Cave *Thiovulum* (see below) and sequences of different *Sulfurimonas* species as an outgroup. A second tree included amplicon sequences as well. For the sake of legibility, the 908 *Thiovulum* sequence variants obtained were clustered at 97 % similarity using CD-HIT-EST (Huang *et al*., 2010), resulting in 50 clusters.

To obtain information on relative *Thiovulum* abundance, the raw short-read libraries (metagenomic) were analyzed with phyloFlash (V 3.3; (Gruber-Vodicka *et al*., 2020)).

### Shotgun sequencing (Illumina and Oxford Nanopore)

Shotgun sequencing was accomplished using both Illumina and Oxford Nanopore sequencing technologies. For Illumina sequencing, 1 ng of genomic DNA from each sample was converted to whole-genome sequencing libraries using the Nextera XT sequencing reagents according to the manufacturer’s instructions (Illumina, San Deigo CA).

A first pass of Oxford Nanopore sequences was obtained using the SQK LSK109 ligation library synthesis reagents on a Rev 9.4 nanopore flow cell with the GridION X5 MK1 sequencing platform, resulting in a total of 131.8 Mbp of reads with a N50 of 1.3 kbp.

Additionally, sequencing was performed on several cellular aggregates that were confirmed microscopically to contain *Thiovulum* cells. The cell aggregates were lysed by freeze thawing and further, following the manufacturer’s instructions, as part of the DNA amplification process using the Repli-G single cell amplification kit (Qiagene, Hilden, Germany). Libraries for Nanopore sequencing were prepared using the LSK-108 kit following the manufacturer’s protocol but skipping the size selection step. The prepared libraries were loaded on MIN106 R9 flow cells, generating a total of 5.7 Gbp of reads with a length N50 of about 3.7 kbp. Basecalling for all Oxford Nanopore reads were done using Guppy 4.0.11.

### cDNA sequencing

cDNA was sheared with the Rapid Shear gDNA shearing kit (Triangle Biotechnology, Durham, NC, USA) and used in the Swift 1S protocol (Accel-NGS 1S Plus kit, Swift Biosciences, Ann Arbor, Mi, USA) with 6 cycles of PCR during indexing. Following library prep, all libraries were pooled in equal volume by combining 2 μl of each library for a final bead clean up with 0.85X AmpPure beads (Beckman Coulter Life Sciences, Indianapolis, IN, USA). This QC pool was then sequenced on an Illumina MiniSeq MO flow cell. The resulting index distribution was used to re-pool the libraries for an Illumina SP flow cell sequencing run with sample LR1 pooled at maximum volume available.

All sequencing data generated in this study were deposited in NCBI Sequence Read Archive under accession number PRJNA673084.

### Metagenomic data analysis

Nanopore reads were assembled using Flye 2.8.1-b1676 (Kolmogorov *et al*., 2019) with default parameters, further manually processed using Bandage (Wick *et al*., 2015) following a final polishing step was performed with unicycler-polish from Unicycler v0.4.9b (Wick *et al*., 2017) using the complete set of Illumina reads (for a total depth of coverage of 12X of the genome) and the subset of Nanopore reads longer than 5 kb (ca. 50X). Polishing consisted of two cycles of pilon 1.23 (Walker *et al*., 2014), one cycle of racon 0.5.0 (Vaser *et al*., 2017) followed by FreeBase (Garrison and Marth, 2012), then 30 additional cycles of short-read polishing using pilon 1.23, after which the assembly reached its best ALE score (Clark *et al*., 2013).

The completeness of the *Thiovulum* genome obtained was assessed using CheckM (Parks *et al*., 2015) and its continuity using the unicycler-check module in Unicycler v0.4.9b. Annotation was performed using Prokka (Seemann, 2014), DRAM (Shaffer *et al*., 2020), KEGG (Kanehisa *et al*., 2016), EggNOG 5.0 (Huerta-Cepas *et al*., 2019), PATRIC (Davis *et al*., 2020; Brettin *et al*., 2015) and RAST (Aziz *et al*., 2008; Overbeek *et al*., 2014). A COG (Tatusov *et al*., 2000) analysis was done using the ANVIO tool (Eren *et al*., 2015). OperonMapper (Taboada *et al*., 2018) was used to inspect the organization of genes into operons. CRISPRs where identified using CRISPR finder tool (Grissa *et al*., 2007). Metabolic models of the annotated genome from Movile Cave and that of *Thiovulum* ES were calculated using PathwayTools (V25.3) (Karp *et al*., 2021).

### Thiovulum sp. genome assembly from public databases

All available metagenomic libraries from the Frasassi caves in Italy (accession numbers in supplementary material) were quality-trimmed using Trimmomatic (Bolger *et al*., 2014) and scanned for the presence of *Thiovulum* 16S rRNA using PhyloFlash (Gruber-Vodicka *et al*., 2020). Library SRR1560850 contained >170,000 *Thiovulum* sp. 16S rRNA and was assembled using Megahit V. 1.2.9 (Li *et al*., 2015), and binned using Metabat2 (Kang *et al*., 2015). The bins were taxonomically annotated using the GTDB-Tk tool (Chaumeil *et al*., 2019) with one bin annotated as *Thiovulum*. The phylogenetic tree generated by the GTDB-Tk tool from a single-copy marker gene multilocus alignment suggested that the Movile and Frasassi caves *Thiovulum* genomes were closely related, hence, both genomes were used to recruit all *Thiovulum* related reads from all Frasassi libraries. The obtained reads were re-assembled, binned and taxonomically annotated, as above, resulting in a *Thiovulum* bin with 94 % completeness, 0.41 % contamination and 25 % strain heterogeneity as evaluated using CheckM (Parks *et al*., 2015).

### Transcriptomic analysis

The 6 libraries containing cDNA sequences (3 from Air Bell 2 and 3 from Lake Room), were quality-trimmed using trimommatic (Bolger *et al*., 2014) and mapped against the complete genome of the Movile cave *Thiovulum* sp. using Salmon version 1.6 (Patro *et al*., 2017). Ribosomal RNA data were removed from the mapping results and TPM (Transcripts Per Kilobase Million) values were recalculated to reflect mRNA expression. The RNA data was analyzed using the iDEP (v. 0.95) online tool (Ge *et al*., 2018) that provides an online graphical user interface for the DeSEQ2 (Love *et al*., 2014) and Limma (Ritchie *et al*., 2015) packages for RNAseq analysis. Differential expression was considered significant with a 2-fold difference and a false discovery rate smaller than 0.1. Taxonomic composition of the active community was obtain by analyzing the 16S rRNA gene from the transcriptomic read libraries using PhyloFlash as above (V 3.3; (Gruber-Vodicka *et al*., 2020)). Viral transcripts were identified using VirSorter2 (v.1.1) (Guo *et al*., 2021), annotated against the viral refseq database release 209 using BLAST and quantified using Salmon version 1.6 (Patro *et al*., 2017).

## RESULTS

### Field observations

A pale-white veil, with a vertical thickness of 2 to 3 mm, was observed below and adjacent to the water surface in Movile Cave (Fig. 2), resembling microbial veils described for sulfur-oxidizing bacteria (Fenchel, 1994; Garcia-Pichel, 1989). Nevertheless, in Movile Cave, the dense agglomeration of cells does not form slime or a strongly cohesive aggregation.

### Microscopy

Veils similar to those seen in the Lake Room (Fig. 2A) were also observed in Air Bell 1 and even more so in Air Bell 2 (Fig 2B) where they reached the highest densities. In Air Bell 2 these veils consisted of large, spherical to ovoid, bacterial cells (Fig. 3A-B) identified as belonging to the genus *Thiovulum*. These cells had a diameter of 12-16 μm, contained 20-30 sulfur globules each (Fig. 3), and occurred in densities of approximately 5.5×10^3^ cells/ml. Transmission Electron Microscope (TEM) observations showed that these large cells were Gram-negative (Fig. 3C-D), and confirmed the existence of 20-30 irregularly shaped sulfur inclusions within each of the cells. Light and TEM imaging revealed *Thiovulum* cell divisions along the long cell axis (Fig. 3B, C). Short peritrichous filamentous appendages (Fig. 3D) observed on the surface of the cells resemble those noticed earlier in other *Thiovulum* species (Wirsen and Jannasch, 1978). Scanning electron microscopy (SEM) revealed the ball-like structure of the sulfur inclusions in a series of connected *Thiovulum* cells (Fig. 3E). These cells were connected one to the other via multiples threads (Fig. 3F-G). Energy-dispersive X-ray (EDX) analysis (Fig. 3H-I) confirmed that the intracellular globules contained sulfur (20.9 - 26.1 %), along with elements common in organic matter such as carbon (49 - 49.2 %) and oxygen (21.1 - 24.6 %), and a few other elements in low abundance such as sodium (2.4 - 3.4 %) and phosphorus (1.2 - 2.2 %).

**Figure 3.**
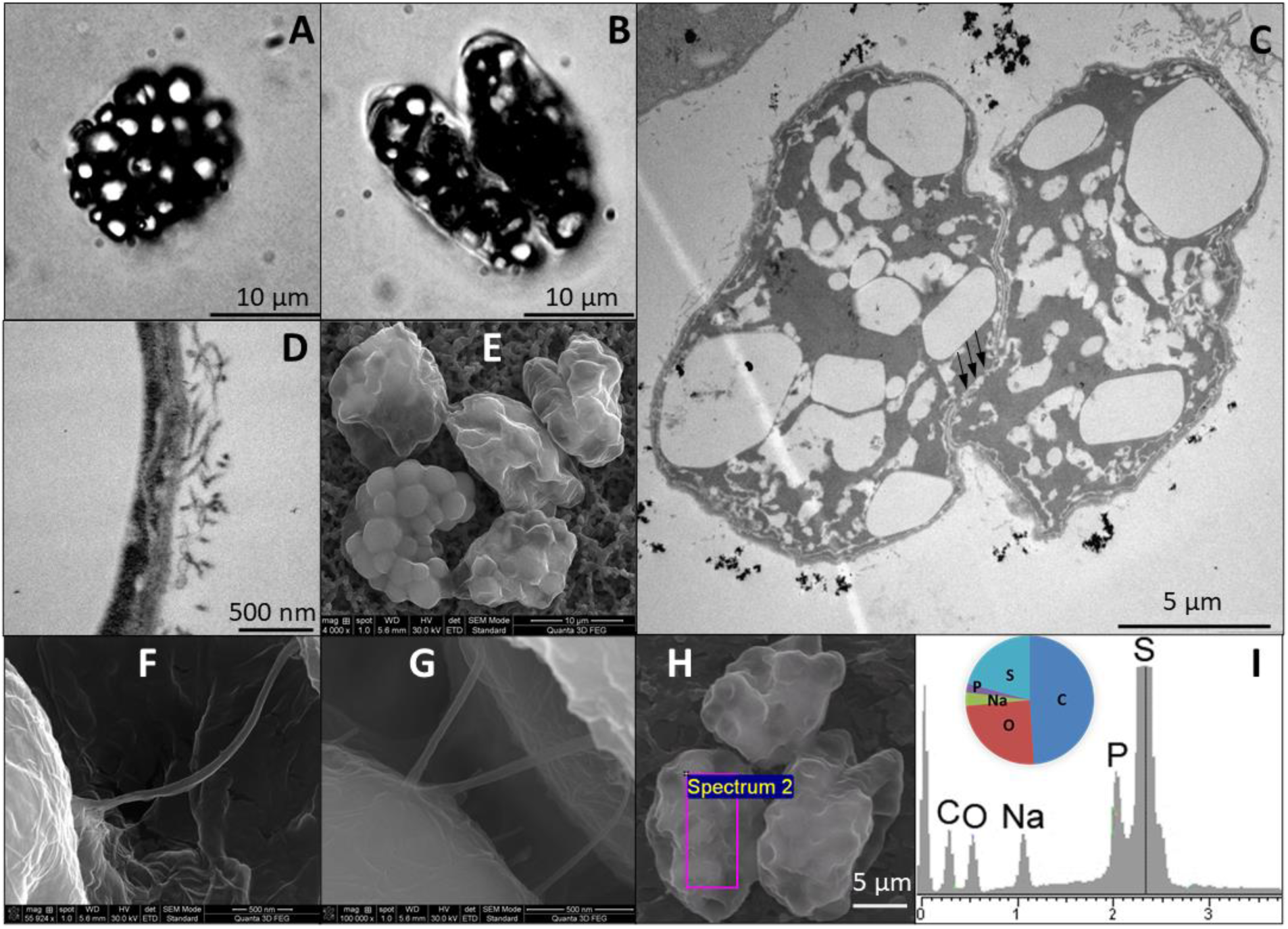
Optical images of giant globular cells colonizing the subsurface veil from Movile Cave (A-B) including a cell undergoing division (B). Each cell carries 20 to 30 sulfur inclusions (large bright spots in panels A, B). TEM images of *Thiovulum* show the cellular localization of sulfur inclusions of various shapes and sizes (panel C, white spots). Ovoid cells divide along their long axis (B, C). The region where the cell membrane is not fully closed between dividing cells is marked with three black arrows (C). The cell wall (D) is covered in pili or short flagella. *Thiovulum* cells (E) are often connected one to another through thread-like structures (F-G). EDX analysis on *Thiovulum* cells (H) inspected under SEM show the typical elemental composition of the cells (I) and confirm the high sulfur content of the internal globules. Note that the height of the peaks in the EDX spectra do not correlate with the element’s ratio but with the X-ray signal intensity.

### Phylogenetic identification and relative abundance of Thiovulum

*Thiovulum* was found in highest abundance (sequence frequency) in Air Bell 2 (35 %), followed by Lake Room (5 %) and submerged microbial mats (0.9 %) (see pie charts in Fig. 1). Similarly, 35 % of the amplicon sequences obtained from Air Bell 2 were annotated as *Thiovulum* sp. A detailed community composition based on 16S rRNA genes obtained from the metagenomic libraries is presented in Supplementary Fig. 1. An amplicon-based multi-year study on the microbial community composition in the cave will be published separately.

The 16S rRNA sequences obtained from the closed genome of the Movile Cave *Thiovulum*, alongside other *Thiovulum* sequences obtained from Movile Cave in an earlier study (Porter *et al*., 2009), formed a separate clade together with other cave and subsurface, freshwater, *Thiovulaceae*, specifically from the sulfidic Frasassi caves in Italy (Fig. 4A). This clade stems from one of two clades of marine *Thiovulaceae*, one of which included *Thiovulum* ES, for which a draft genome is available (Marshall *et al*., 2012). Including shorter amplicon sequences of *Thiovulum* from Air Bell 2 in the phylogenetic tree (Supplementary Fig. 2), highlights the diversity of these bacteria in the cave.

**Figure 4.**
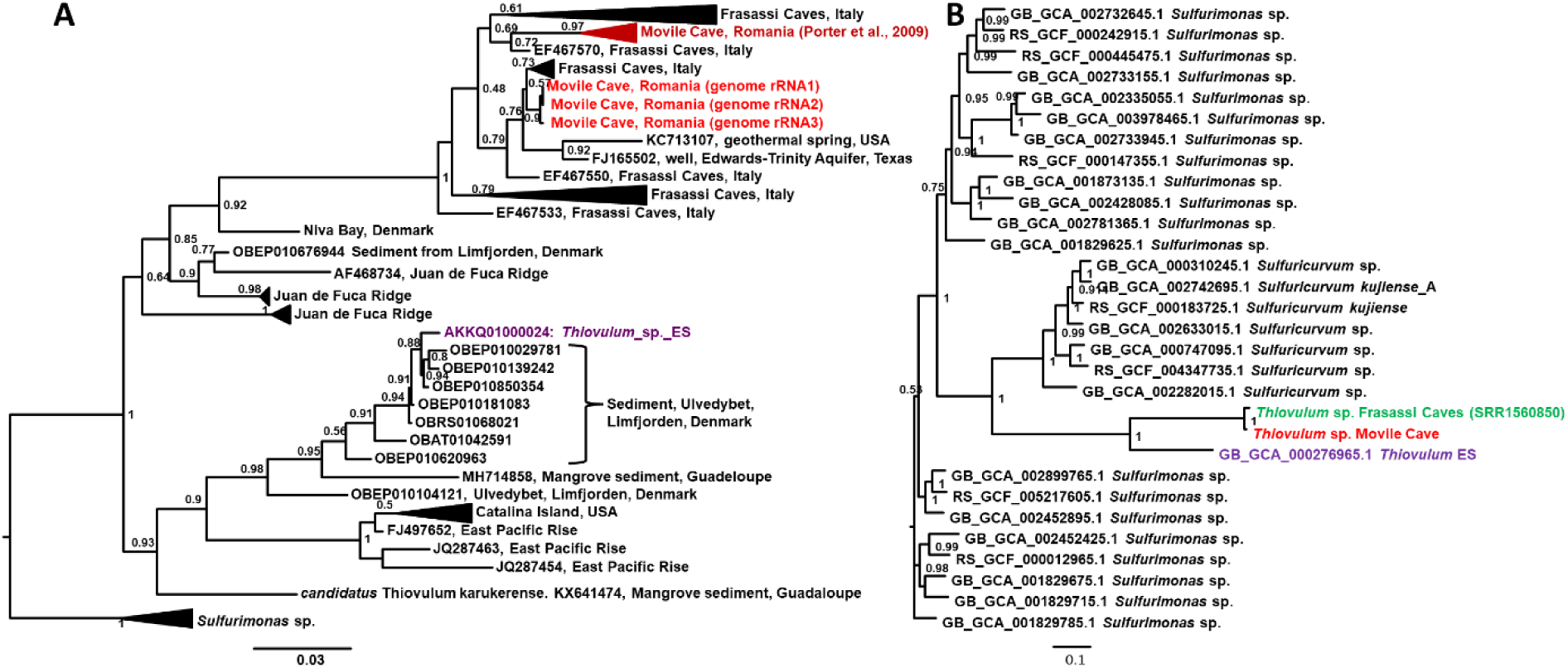
Maximum-likelihood placement of the 16S rRNA gene of *Thiovulum* from Movile Cave (A) and its genome (B). The 16S rRNA sequences obtained from the complete genome of the Movile Cave *Thiovulum* are shown besides all *Thiovulum* sequences available in the SILVA database (V138.1 Quast et al 2013) using *Sulfurimonas* sp. as an outgroup. A similar 16S rRNA tree using also *Thiovulum* amplicon sequences from Movile Cave is shown in Fig. S2. The multilocus protein alignment, of single-copy marker genes from the Movile Cave *Thiovulum*, is shown together with that obtained from the genome of *Thiovulum* ES and a *Thiovulum*-annotated bin from public metagenomic data from the Frasassi caves (SRR1560850). The protein alignment was generated using GTDB-TK (Chaumeil *et al*., 2019). The Shimodaira-Hasegawa local support values (ranging from 0 to 1) are shown next to each node.

A phylogenetic tree constructed from a multilocus alignment of single-copy marker genes from *Thiovulum* ES (Marshall *et al*., 2012), the *Thiovulum* genome from Movile Cave, a metagenome assembled genome from the Frasassi caves, and all available *Sulfurimonas* genomes, resulted in the Frasassi and Movile *Thiovulum* genomes forming a separate clade (Fig. 4B). The Movile Cave genome had an overall low similarity to the marine *Thiovulum* (ES) genome (Marshall *et al*., 2012), with an average nucleotide identity (ANI) of 74.49 %, an average amino acid identity (AAI) of 58.33 % (Fig. 5A-B), and a very low sequence synteny (Fig. 5C). In contrast, The Movile and Frasassi *Thiovulum* genomes were highly similar with an ANI of 97 % and an AAI of 95 %. (Fig. 5D, E), as well as a high gene synteny (Fig. 5F).

**Figure 5.**
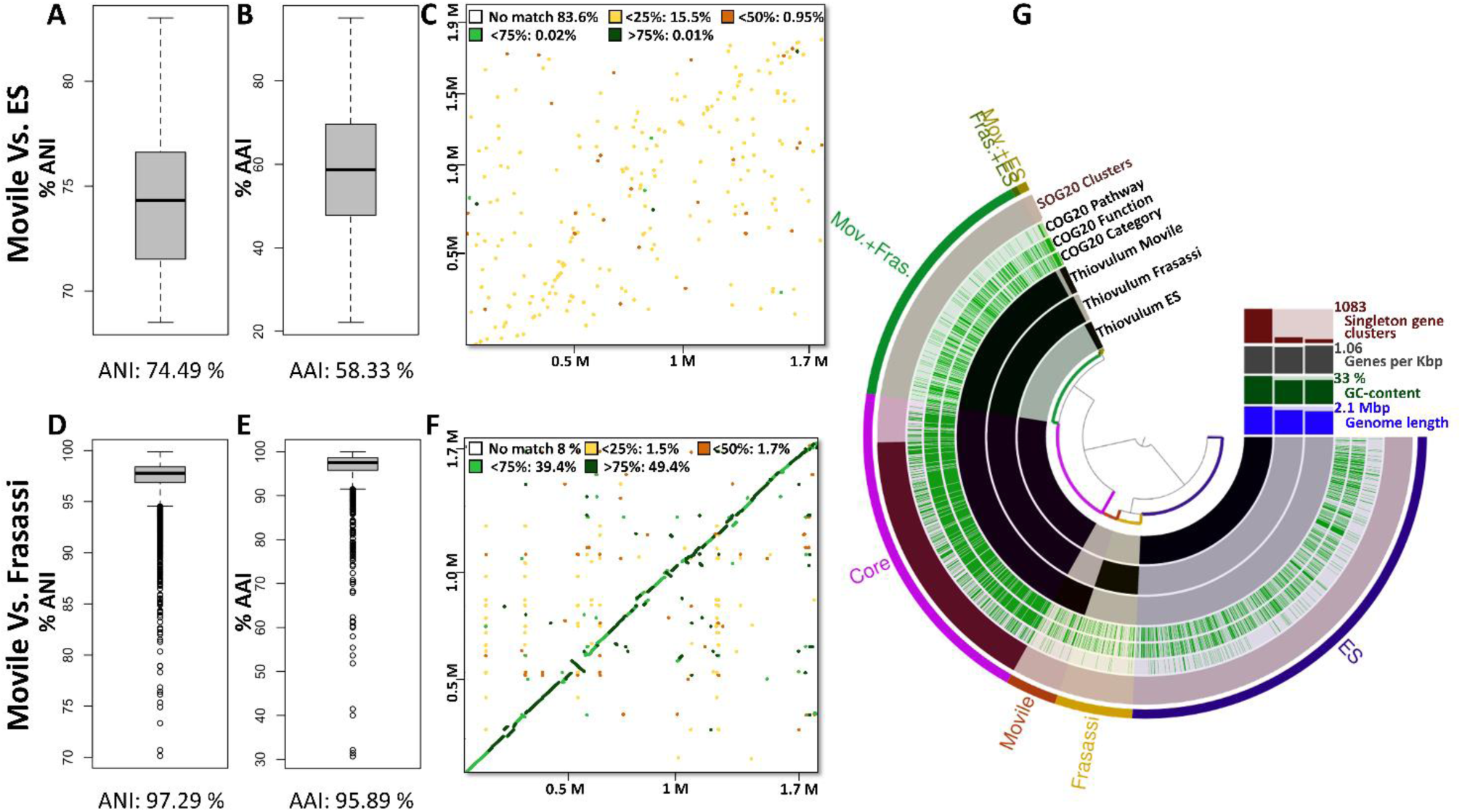
The genome of the Movile *Thiovulum* strain, compared to *Thiovulum* ES (Marshall *et al*., 2012) (A-C) and to the Frasassi *Thiovulum* (D-F), using Average Nucleotide Identity (ANI) (A,D), Average Amino acid Identity (AAI) (B,E), contig mapping against the genome of the Movile Cave *Thiovulum* (C,F) and COG annotation (G).

### Genome analysis

The assembly of metagenomic data from Movile Cave resulted in a closed circular genomic sequence classified as *Thiovulum* sp. with a genome length of 1.75 Mbp (coverage X330) and a GC content of 28.4 %. Genome completeness was estimated using CheckM (Parks *et al*., 2015) at 93 %. Using a *Campylobacteraceae* specific set of marker genes did not improve the completeness prediction, however, in absence of sufficient reference genomes for this genus, this value likely represents the full set of marker genes for *Thiovulum*. CheckM estimated a contamination of 0 % and a strain heterogeneity of 0, suggesting the *Thiovulum* genome assembly does not contain any contaminating sequences from additional distant or closely related organisms.

The genome was analyzed using different tools with the results summarized in Supplementary Dataset 1. Genes discussed further on are addressed using the notation G2Y-n, where n refers to an incremental number. This notation is used by PathwayTools (Karp *et al*., 2021) and match the supplementary metabolic model provided (Supplementary Figures 3,4).

The genome contains 1804 coding sequences, of which 1534 could be annotated and 270 remain hypothetical proteins, 36 tRNAs genes, 3 rRNA operons and 9 CRISPR arrays in which 5 Type III Cas genes were identified (G2Y-562:567), comprising a total of 77 repeats. The same annotation conducted de novo on the *Thiovulum* ES genomes suggests, based on COGs (Clusters of Orthologs genes), that the Movile, ES and Frasassi strains share 879 core genes (Fig. 5G and Supplementary Dataset 2). The Movile strain further shares 33 and 777 genes with the ES and Frasassi strains, respectively. The Frasassi and ES strains had 26 common genes in addition to the core genome. The Movile, ES, and Frasassi genomes further contained 145, 1201, and 207 individual genes, respectively.

### Carbon metabolism

Similar to *Thiovulum* ES, all genes required for C fixation via the reductive TCA cycle could be identified. in the *Thiovulum* genomes from Movile Cave and from Frasassi. The oxidative TCA cycle is complete as well in both the *Thiovulum* species with the citrate synthase gene (EC 2.3.3.1) replaced by ATP-citrate lyase (EC 2.3.3.8; GDY-1367,1367 alpha and beta subunits, respectively) (Fig. 6, Supplementary Figs 3,4, Supplementary Dataset 1).

**Figure 6.**
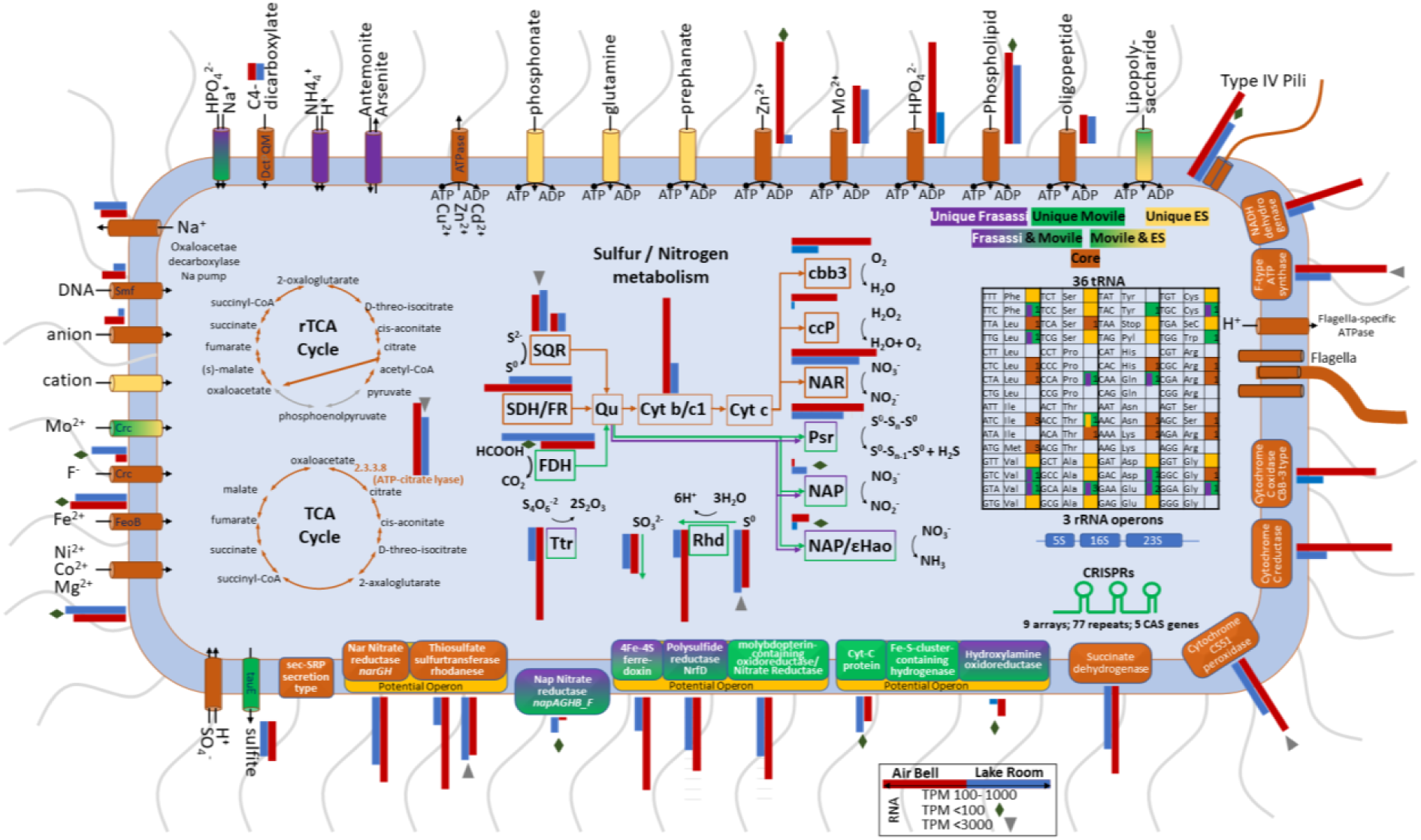
Graphical summary of main components of the Movile, ES, and Frasassi *Thiovulum* strains. Elements or reactions colored in green, orange or purple are unique to the species from Movile, ES or Frasassi, respectively. Core elements or reactions are colored in brown. Gradient colors indicate presence in two of the genomes. Grey arrows in the reductive TCA (rTCA) cycle show missing reactions. The sulfur/nitrogen metabolism model was drawn based on Grote *et al*. (2012), Hamilton *et al*. (2014), and Poser *et al*. (2014). SQR: sulfide-quinone oxidoreductase, SDH/FR: succinate dehydrogenase/fumarate reductase, FDH: formate dehydrogenase, Qu: quinone, Cyt b/c1, quinone cytochrome oxidoreductase, cbb3, cytochrome c oxidase; ccp: cytochrome c peroxidase, NAP, periplasmic nitrate reductase; NAR, membrane bound nitrate reductase; Psr, polysulfide reductase, εHao: Epsilonproteobacterial hydroxylamine oxidoreductase, Ttr: tetrathionate reductase, Rhd: rhodanese-related sulfurtransferase. CRISPR were not identified in the genome of *Thiovulum* ES, probably due to the current fragmented nature of the data. Comparative gene expression between *Thiovulum* in Air Bell 2 (red) and the Lake Room (blue) are shown for each gene depicted where the TPM value was above 10. Genes were the TPM value was below 100 of above 1000 are marked with a diamond and inverse triangle, respectively. For proteins consists of multiple subunits, the expression is the genes encoding for one of the subunits. For SQR and RhD expression is show for both copies of the genes.

### Sulfur metabolism

All annotation approaches (Supplementary Dataset 1) revealed only few genes involved in dissimilatory sulfur cycling, including two copies of the sulfide:quinone oxidoredutase (G2Y-583, G2Y-1704) that oxidizes sulfide to polysulfide, and the polysulfide reductase gene *nrf*D (G2Y-67) that carries out the reverse process. *nrf*D was found in a 3-gene potential operon together with the large subunit of the assimilatory nitrate reductase (*nar*B, G2Y-68) and the *ttr*B; tetrathionate reductase subunit B (G2Y-66). Two rhodanese sulfur transferase proteins (G2Y-815, G2Y-816) were identified in a 6-gene operon containing two other subunits of a nitrate reductase (*nar*H: G2Y-813 and *nar*G: G2Y-814). Among the two other genes in this operon, one is related to cytochrome C (G2Y-811) and the other could not be annotated (G2Y-810). The *tau*E sulfite exporter was identified (G2Y-644) as part of a 5-gene operon involved in the transport of molybdate (G2Y-641-643,645 *mod*CABD, respectively). Sulfite dehydrogenase, dissimilatory sulfite reductase (*dsr*AB), the sox genes or adenylyl sulfate reductase (*apr*AB) that carry out the sulfide oxidation to SO _4_^2-^ could not be found by any of the annotation tools nor by manual BLAST against all sequences available for each of those protein in the UniProt database.

### Nitrogen metabolism

In addition to the membrane-bound nitrate reductases (*nar*, G2Y-813,814) found also in *Thiovulum* ES, the Movile and Frasassi cave *Thiovulum* possesses also periplasmatic nitrate reductases encoded by the *nap* genes encoded in one operon (G2Y-1099-G2Y-1099, *nap*AGHB_F). The hypothetical protein encoded in this operon (G2Y-1098) is likely part of the *nap*F gene (G2Y-1099) as seen by BLAST analysis in other *Campylobacteraceae*. Additionally, the gene for hydroxylamine dehydrogenase, which is often encountered in genomes from *Campylobacterota* (Haase *et al*., 2017), formerly referred to as *Epsilonproteobacteria* (Waite *et al*., 2019), was also identified (G2Y-1392) in a 3-gene operon with two unannotated hypothetical genes (G2Y-1390, G2Y-1391).

### Chemotaxis and motility

As *Thiovulum* sp. is a highly motile bacterium, we inspected motility and chemotaxis genes. All genes necessary for flagellar assembly were found in the Movile and Frasassi strains, similarly to *Thiovulum* ES. The chemotaxis genes *che*V, *che*A, *che*W, *che*D (G2Y-470:472 *che*VAW, G2Y-741 *Che*D) and *che*Y (G2Y-1156) were identified as well as additional *che*Y-like domains (G2Y-6,20,151,182,251,389,582,712,973,1116,1130,1156’,1344,1486,1516). The *cet*A and *cet*B (G2Y-174:176 *cet*ABB’) energy taxis genes and the parallel to the *Escherichia coli* aerotaxis (*aer*) (G2Y-1557) gene were also identified. The *che*X gene, which was not found in the genome of *Thiovulum* ES, was identified in the Movile Cave *Thiovulum*. Gene *che*B was reported missing in *Thiovulum* ES, was identified in the Movile strain (G2Y-1843) but also in *Thiovulum* ES upon COG reannotation. Additionally, 9 methyl-accepting chemotaxis proteins were identified (G2Y-84,85,181,721,740,1225,1487,1557,1836).

In addition to flagella genes, (G2Y-3, *fli*C; G2Y-45, *flg*A; G2Y-48, *fli*C; G2Y-184, *flh*B; G2Y-302, *mot*B; G2Y-336, *flg*K; G2Y-338, *flg*M; G2Y-350, *fli*N; G2Y-367, *fli*F; G2Y-442, *flh*A; G2Y-569, *flg*H; G2Y-656, *fli*G; G2Y-666, *lag*; G2Y-781, *fli*N; G2Y-790, *fli*I; G2Y-928, *flg*E; G2Y-1052, *fli*S; G2Y-1053, *fli*D; G2Y-1054, *lag*; G2Y-1107, *flg*E; G2Y-1108, *flg*D; G2Y-1122, *flg*B; G2Y-1199, *fli*H; G2Y-1218, *fli*M; G2Y-1222, *flh*F; G2Y-1250, *fli*R; G2Y-1258, *flg*I; G2Y-1331, *fli*L; G2Y-1458, *mot*A; G2Y-1522, *flg*G; G2Y-1568, *fli*Q; G2Y-1728, *flg*F; G2Y-1801, *fli*E; G2Y-1802, *flg*C) the *pil*A, *pil*E, *pil*T, *pil*N, *pul*O, *fim*V genes responsible for the formation and retraction of type IV pili were identified.

### Gene expression

Samples from Air Bell 2 and from the Lake Room were collected for RNA analysis to confirm that *Thiovulum* is active in Movile Cave. 16S rRNA of *Thiovulum* dominated all samples, making up more than 94 % of the active community (Figure S1) even though *Thiovulum* DNA was rare in the Lake Room in our previous samples. Despite the similarity in abundance, the gene expression profiles differ significantly between the two sites (Fig. 7). In both the heatmap (Fig. 7A) and the principal component analysis (Fig. 7B), the samples from the different environments clustered separately, with clear clusters of genes differently expressed in the two cave compartments. Differential expression analysis (Fig. 7C; Supplementary Dataset 3) revealed that 222 genes were more expressed in the Lake Room compared to Air Bell 2, while the opposite comparison resulted in 42 genes. Over half of the genes more expressed in the Lake Room encoded for hypothetical proteins to which no function could be assigned. Retron-type reverse transcriptases were the most dominant group of genes (n=15) also exhibiting some of the highest transcription level with TPM values up to 19,000. Genes over expressed in samples from Air Bell 2 were related to energy generation including cytochromes c and b as well as F-type ATP synthase. The entire gene expression data is available in Supplementary Dataset 1 and is additionally depicted in Fig. 6 next to the displayed genes or functions.

**Figure 7.**
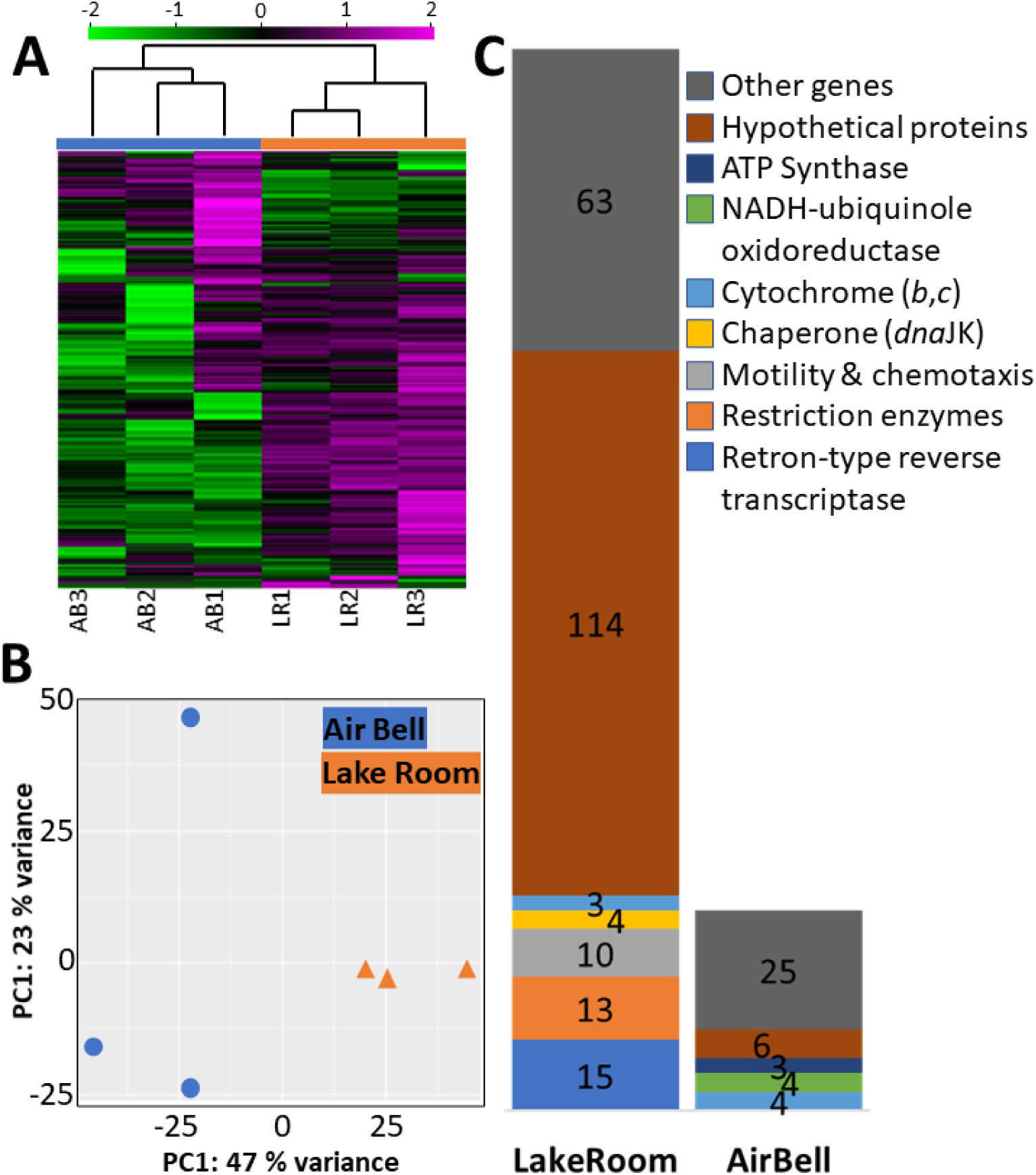
Comparison of mRNA transcriptomic profiles of the Movile Cave *Thiovulum* obtained from triplicates samples collected in Air Bell 2 and the Lake Room. A heatmap shows that the two environments are separated from one another with clusters of genes expressed more or less in one of the two environments (A). The values in the heatmap are log-transformed TPM values and normalized using each gene’s standard deviation. Principle components analysis (B) demonstrates the separation between samples mainly across PC1 likely representing sample location. Differential expression analysis (C) revealed that 222 genes are significantly more expressed in the Lake Room as compared to Air Bell 2, whereas 48 genes. are significantly less expressed

## DISCUSSION

In Movile Cave, the oxidation of reduced compounds such as H_2_S, CH_4_, and NH_4_^+^ is the only primary energy source. *Thiovulum*, a large sulfur oxidizer, often found in close proximity to sediments, microbial mats or surfaces (Marshall *et al*., 2012; Jorgensen and Revsbech, 1983; Gros, 2017), is part of the Movile chemoautotrophic microbial community, involved in *in situ* carbon fixation, that represents the base of the cave’s food web that supports an abundant and diverse invertebrate community (Sarbu, 2000; Brad *et al*., 2021). These *Thiovulum* cells, exceeding 15 μm in diameter, are larger than most known sulfur-oxidizers, belonging to the group of giant sulfur bacteria (Ionescu and Bizic, 2019). Here we investigated the morphological, phylogenetic, and genomic aspects of a fully planktonic *Thiovulum* sp.. We further compared its genome to that of the sole other existing genome of *Thiovulum*, strain ES. The latter, originating from a phototrophic marine mat, was reannotated for the purpose of this comparison to account for new available information, 8 years after its original publication. The main aspects of this comparison are depicted in Fig. 6 and in more details in Supplementary Fig. 3 and 4. The close phylogenetic relationship of the Movile Cave *Thiovulum* 16S rRNA gene with sequences from the Frasassi caves prompted us to recover a *Thiovulum* genome from publicly available data. The obtained MAG was added to the discussed comparison.

### Hypoxic Air Bell 2 vs. oxic Lake Room

*Thiovulum* is not typically dominant in microscopy-or DNA based observations from the Lake Room. Yet, its rRNA gene dominated over 94 % of the transcriptomic data, similar to its presence in the RNA samples from the hypoxic Air Bell 2. Nevertheless, it is known that community profiles obtained from DNA representing pseudo-abundance, and those from RNA, representing degree of activity, can substantially differ one from the other (e.g. Shu *et al*., 2019; Bižić-Ionescu *et al*., 2018). The presence and high activity of *Thiovulum* at the surface of both the Lake Room and Air Bell 2, environments that differ significantly in the overlaying atmosphere, points to the metabolic flexibility of cave-dwelling *Thiovulum* strains and perhaps of the entire genus.

While the two expression profiles differed significantly, it is evident (Fig. 6) that most genes recognizable as involved in cell metabolism had higher expression levels in Air Bell 2, though not all at statistically significant levels (Supplementary Dataset 1). More than half of the genes overexpressed by *Thiovulum* in the Lake Room could not be assigned any function making it impossible to understand it’s specific metabolic activity in that compartment of the cave. However the high expression of retron-type reverse transcriptase and Type II restriction enzymes in the Lake Room can be indicative of an ongoing phage infection (Millman *et al*., 2020; Pingoud *et al*., 2014), which may explain the reduced metabolic activity and elevated expression of defense systems, though only one CRISPR associate gene was overexpressed. Quantification of viral transcripts showed an overall higher expression in the Lake Room (Fig. S6), however, at this stage it is not possible to directly connect these transcripts to *Thiovulum*.

### Phylogeny

Our phylogenetic analysis of all available *Thiovulum* spp. 16S rRNA sequences revealed two main clades of marine origin with no clear physical or biogeochemical basis for the separation. All sequences obtained from sulfidic caves, or subsurface environments (e.g., a drinking water well) formed a subclade in one of these two clades. This evolutionary transition from a marine environment towards a freshwater one likely accounts for the differences in sequence and function between *Thiovulum* ES, found in a phototrophic mat in a marine environment, and the Movile and Frasassi cave *Thiovulum*. This hypothesis should be further validated as more genomes of *Thiovulum* will become available.

### Sulfur and nitrogen metabolism

*Thiovulum* presents several interesting features, such as being one of the fastest bacterial swimmers and being able to form large veils consisting of interconnected cells. Many sulfur-oxidizing microorganisms including species of *Thiovulum* form veils by means of what appear to be mucous threads. These threads are used by the cells to attach to solid surfaces (Fauré-Fremiet and Rouiller, 1958; Fenchel, 1994; Thar and Fenchel, 2001; De Boer *et al*., 1961; Wirsen and Jannasch, 1978; Robertson *et al*., 2015). In marine settings, such veils keep cells above sediments (Karavaiko *et al*., 2006) at the oxic-anoxic interface where the optimal concentration of O_2_ and H_2_S can be found.

SEM analyses indicated that the cells in the dense agglomeration in Movile Cave are at least partially interconnected. It has been hypothesized that the coordinated movement of *Thiovulum* cells generates convective transport of H_2_S or O_2_ to the cells (Petroff *et al*., 2015; Fenchel and Glud, 1998). The fully planktonic localization of the cells in Air Bell 2 means that *Thiovulum* here cannot use surfaces to place itself at the oxic-anoxic interphase. O_2_ in the Lake Room was shown to penetrate only the upper 1 mm of the water (Riess *et al*., 1999), and this is likely similar in the hypoxic Air Bell 2.

*Thiovulum* is a sulfide oxidizer as evidenced by the generation and accumulation of sulfur inclusions. The amount and type of sulfur inclusions in cells is influenced by the concentrations of H_2_S and O_2_ in the environment. Typically, cells store elemental sulfur when H_2_S is abundant in the environment, and later use the intracellular reserves of sulfur when the sulfide source in the environment is depleted (De Boer *et al*., 1961). Sulfur inclusions were also shown to form when the supply of O_2_ is limited and as a result the sulfur cannot be entirely oxidized to soluble sulfite, thiosulfate, or sulfate. Complete depletion of sulfur inclusions from cells is not likely in Movile Cave where abundant H_2_S is available (245 μM (Flot *et al*., 2014)) continuously and where O_2_ is scarce in most habitats, and specifically in Air Bell 2 (Sarbu *et al*., 1996). The analysis of the Movile Cave, ES, and Frasassi strains genomes identified the SQR gene responsible for the oxidation of sulfide to elemental sulfur. Nevertheless, the genes required for further oxidizing elemental sulfur to sulfate, via either of the known mechanisms, were not found. An exception to this is the possible oxidation of sulfite to sulfate via the intermediate adenylyl sulfate by *Thiovulum* ES, for which the gene encoding the sulfate adenylyl transferase was originally found (Marshall *et al*., 2012), yet, according to our re-annotation the necessary adenylyl-sulfate reductase genes *apr* (EC1.8.4.9) or *apr*A (EC1.8.99.2) are missing.

Marshall *et al*. (2012) proposed that *Thiovulum* undergoes frequent (daily) oxic/anoxic cycles that prevent continuous accumulation of elemental sulfur in the cell. We advance three additional options by which the Movile Cave and likely the Frasassi caves *Thiovulum*, may avoid sulfur accumulation. First, the presence of a polysulfide reductase (*nrf*D) suggests that the cells can reduce polysulfide back to sulfide (Fig. 6). Second, the identification of different rhodanese genes, known to be involved in thiosulfate and S^0^ conversion to sulfite (Poser *et al*., 2014), and of a sulfite exporter (*tau*E) in the Movile Cave strain, suggests that *Thiovulum* may be able oxidize elemental sulfur to sulfite and transport the latter out of the cell. Third, we propose that cave-dwelling *Thiovulaceae* are capable of dissimilatory nitrate reduction to ammonia (DNRA) using elemental sulfur (Slobodkina *et al*., 2017), a process already shown in *Campylobacterota* (e.g. *Sulfurospirillum deleyianum*) (Eisenmann *et al*., 1995). The Movile and Frasassi *Thiovulum* contain not only the *nar* (*nar*GH) genes for nitrate reduction, but also the periplasmatic *nap* genes known for their higher affinity and ability to function in low nitrate concentrations (Pandey *et al*., 2020). Additionally, they harbor the gene for the epsilonproteobacterial hydroxylamine dehydrogenase (ε*hao*). Hydroxylamine dehydrogenase is known from other *Campylobacterota* (e.g. *Campylobacter fetus* or *Nautilia profundicola*) and was shown to mediate the respiratory reduction of nitrite to ammonia (Haase *et al*., 2017). In line with the findings of Marshall *et al*. (2012), the *hao* gene was not found in the genome of *Thiovulum* ES upon re-annotation, suggesting that the *hao* gene may not be part of the core *Thiovulum* genome. Normally, *Campylobacterota* that utilize hydroxylamine dehydrogenase do not have formate-dependent nitrite reductase, matching the annotation of the Movile Cave *Thiovulum. Campylobacterota* typically use periplasmic nitrate reductase (*nap*) and do not have membrane-bound *nar*GHI system (Kern and Simon, 2009; Meyer and Huber, 2014). Interestingly, *Thiovulum* ES has only Nar systems while the Movile and Frasassi strains have both types, suggesting that *nap* genes may be a later acquisition by cave-dwelling *Thiovulaceae*. Nevertheless, while genomic information is suggestive of the presence or absence of specific enzymes and pathways, additional experiments and gene expression data are required to determine which of the genes are utilized and under which environmental conditions.

Thus, we propose that, if O_2_ is available, sulfide is oxidized to elemental sulfur with oxygen as an electron acceptor (as may have been the case for part of the community at the time of sampling give the high expression of cytochrome *c* oxidase cbb3; Fig. 6). However, when cells are located below the O_2_ penetration depth, the Movile Cave *Thiovulum* may oxidize sulfide using NO_3_^-^ as an electron acceptor, in a process of dissimilatory nitrate reduction to ammonium, as documented in other *Campylobacteraceae*. Our transcriptomic analysis, however, point out that at the time of sampling the *nap* and ε*hao* genes were minimally expressed as compared to other sulfur and nitrogen metabolism genes (Fig. 6, Supplementary Dataset 1). Even though ε*hao* expression was more than 3 times higher in samples from Air Bell 2 than in Lake Room, this suggests that the DNRA pathway was not highly active in the *Thiovulum* community. In contrast, the high expression of both copies of rhodanase genes as well sulfite exporter (*tau*E) suggest that elemental sulfur may have been converted to sulfite and excreted.

### Cell motility and veil formation

*Thiovulum* sp. often forms large veils of interconnected cells. The threads connecting the cells are thought to be secreted by the antapical organelle located at the posterior side of the cell (De Boer *et al*., 1961; Robertson *et al*., 2015). Short peritrichous filaments (Fig. 3D) observed on the surface of the cells from Movile Cave resemble those noticed earlier in *Thiovulum* species and referred to as flagella (Wirsen and Jannasch, 1978). While all genes necessary for flagella assembly were found in the Movile Cave, Frasassi and ES *Thiovulum* strains, so were genes for type IV pili. Evaluating available electron microscopy images, we suggest that these ideas need to be revisited.

Our SEM images (as well as previous ones of connected *Thiovulum* cells) show connecting threads that are not exclusively polar and are much thinner than the stalk-like structure shown by de Boer *et al*. (De Boer *et al*., 1961). We propose that these structures are rather type IV pili, which are known, among other functions, to connect cells to surfaces or other cells (Craig *et al*., 2019). Alternatively, Bhattacharya *et al*. (2019) have shown the formation of cell-connecting nanotubes constructed using the same enzymatic machinery used for flagella assembly. However, it is not possible to determine this in absence of TEM images of connected cells showing the presence of the reduced flagellar base.

We further question the flagellar nature of the peritrichous structures around the cells. Inspecting the high-resolution electron microscopy images taken by Fauré-Fremiet and Rouiller (1958), there is no single structure visible resembling a flagellar motor. Additionally, the length of these structures based on de Boer *et al*. (1961) and a similar image in Robertson *et al*. (2015) suggest that these < 3 μm structures are too short for typical flagella (> 10 μm in length, c.f. (Renault *et al*., 2017)) and are closer to the 1-2 μm lengths known type for pili. Interestingly, *Caulobacter crescentus* swims at speeds of up to 100 μm s^-1^ using a single flagellum aided by multiple pili (Gao *et al*., 2014). In light of this hypothesis, we inspected the images of the fibrillar organelle at the antapical pole of the cells. The high-resolution images presented in Fauré-Fremiet and Rouiller (1958) and in de Boer (De Boer *et al*., 1961) show an area of densely packed fibrillar structures. Considering our current knowledge in flagellar motor size (ca. 20 nm) it is highly unlikely that each of these fibres is an individual flagellum, thus, potentially representing a new flagellar organization. Furthermore, none of our TEM images could reveal flagella motor-like structures on the cell. Given the high number of flagella-like structure around the cell it is logical to assume that at least some would be seen in the images taken. Petroff *et al*. (Petroff *et al*., 2015) investigated the physics behind the 2-dimensional plane assembly of *Thiovulum* veils and suggested it to be a direct result from the rotational movement attracting cells to each other. Nevertheless, as seen, SEM images show cells that are physically attached one to the other, suggesting several mechanisms and steps may be involved. Interestingly, type IV pili retraction can generate forces up to 150 pN known to be involved in twitching motility in bacteria (Craig *et al*., 2019). If coordinated, these may be part of the explanation of the swimming velocity of *Thiovulum* which is, at ca. 615 μm s^-1^, 5 to 10 times higher than that of other flagellated bacteria (Garcia-Pichel, 1989). Thus, genomic information and re-evaluation of electron microscopy data raise new questions concerning the nature of the extracellular structures on the surface of *Thiovulum* sp. and call for new targeted investigations into this topic.

## CONCLUSIONS

Movile Cave is a ecosystem entirely depending on chemosynthesis. We showed that submerged near-surface, planktonic microbial accumulations are dominated by *Thiovulum*, a giant bacterium typically associated with photosynthetic microbial mats. We further showed that *Thiovulum* dominates the active fraction in surface waters of hypoxic and oxic compartments of the cave, suggesting metabolic flexibility

Our results highlight the existence of a clade of cave and subsurface *Thiovulaceae* that based on genomic information differs significantly from marine *Thiovulum*. The genomes of this planktonic *Thiovulum* strain as well as that of the highly similar *Thiovulum* from the Frasassi sulfidic caves suggest that these can perform dissimilatory reduction of nitrate to ammonium, when O_2_ is unavailable. Thus, *Thiovulum* may play a role in the nitrogen cycle of sulfidic caves, providing readily available ammonia to the surrounding microorganisms. The coupling of DNRA to sulfide oxidation provides a direct and more productive source of ammonium.

This investigation of three *Thiovulum* genomes, coupled with observations of current and previous microscopy images, questions the number of flagella the cells have, bringing forth the possibility that the cells may use type IV pili for rapid movement and cell-to-cell interactions.

The collective behavior of *Thiovulum* is still a puzzle and there may be more than one mechanism keeping the cells connected in clusters or in veils. Our SEM images suggest the cells are connected by thread-like structures. Petroff *et al*. (2015) show, on another strain, that there is no physical connection between the cells. They suggest that the swimming behavior of individual cells keeps the cells together. More research is therefore needed to understand if these different mechanisms are driven by strain variability, or by different environmental conditions.

## Supporting information

Supplementary_data1_Merged_annotation_with_RNASeq_Thiovulum_Movile

Supplementary_data2_COG_Summary_Movile_ES_Frasassi

Supplementary_data3_Overall_Gene_expression

## ACKNOWLEDGEMENTS

The authors thank GESS team for logistics with sampling in the cave. We would also like to thank Pheobe Laaguiby at the University of Vermont for performing the outstanding Oxford Nanopore sequencing and Bo Barker Jørgensen, Emily Fleming, Carl Wirsen, and Tom Fenchel for their valuable suggestions that led to the improvement of the quality of this manuscript. We would also like to thank the Extreme Microbiome Project (XMP) for providing the DNA extraction reagents and methods as well as Laura Gray and Mehdi Keddache at Illumina Corp for providing partial sequencing reagents through its partnership with XMP. T. Brad was supported by a grant of Ministry of Research and Innovation, project number PN-III-P4-ID-PCCF-2016-0016 (DARKFOOD), and by EEA Grants 2014-2021, under Project contract no. 4/2019 (GROUNDWATERISK). S. M. Sarbu was supported by a grant of Ministry of Research and Innovation (UEFISCDI) projects number PN-III-P4-ID-PCE-2020-2843 (EVO-DEVO-CAVE). J.W. Aerts acknowledges the support from a grant from the User Support Programme Space Research (grant ALW-GO/13-09) of the Netherlands Organization for Scientific Research (NWO). M. Bizic was funded through the German Research Foundation (DFG) Eigene Stelle project BI 1987/2-1. The computational resources for the assembly of the *Thiovulum* genome were provided to J.-F. Flot by the Consortium des Équipements de Calcul Intensif (CÉCI) funded by the Fonds de la Recherche Scientifique de Belgique (F.R.S.-FNRS) under Grant No. 2.5020.11; D. Ionescu was funded through the German Research Foundation (DFG) Eigene Stelle project IO 98/3-1.

