## Supplementary text and figures for "Cave *Thiovulaceae* differ metabolically and genomically from marine species"

Supplementary Material

### DETAILED MATERIALS AND METHODS

Replicate samples of water were collected into sterile containers from the surface of the small sulfidic lake and from the Air Bells in the lower section of Movile Cave (Fig. 1). Unpreserved 50 ml water samples were immediately brought to the laboratory and inspected using optical microscopy, while other 50 ml water samples were preserved with ethanol to a final concentration of 50 % and later used for DNA / RNA extraction. Additionally, 15-ml water samples were preserved with formaldehyde to a final concentration of 4 % and were later used for cell enumeration. Samples used for DNA extraction were collected in July 2019 whereas samples for RNA extraction were collected in August 2021.

#### *Electron microscopy and elemental analysis*

Two 15-ml samples collected from the water surface in Movile Cave were centrifuged, and the microorganisms were resuspended and fixed for 2 h with 2.7 % glutaraldehyde in phosphate buffered saline (1× PBS). The cells were then rinsed three times in 1× PBS, and finally fixed for 1 h with 2 % osmic acid in 1× PBS. The cells were harvested again by centrifugation, dehydrated in graded acetone-distilled water dilutions, and embedded in epoxy resin. Sections of about 100 nm thickness were produced with a diamond knife (Diatome, Hatfield) using a Leica UC6 ultramicrotome (Leica Microsystems, Wetzlar, Germany) and were stained with lead citrate and uranyl acetate (Hayat, 2001). The grids were examined with a Jeol JEM transmission electron microscope. Samples for scanning electron microscopy (SEM) were fixed with 2.7 % glutaraldehyde in 1× PBS, air dried on 0.22 µm mesh-sized Millipore filters, sputter coated with 10 nm gold and examined on a JEOL JSM 5510 LV microscope (Jeol, Japan). Energy-dispersive X-ray spectroscopy (EDX) analysis was performed with an EDX analyzer (Oxford Instruments, Abingdon, UK) and with the INCA 300 software.

#### *gDNA Extraction*

Bacteria from water samples were concentrated using a vacuum pump and Nalgene single-use analytical filter funnels (Thermo Fisher Scientific, MA, USA), with the included filter replaced with a 0.2 µm isopore membrane filter (Millipore Sigma, MA, USA). Prior to filtration, the glass assembly components were autoclaved at 121.5 °C for 30 min wet and 20 min dry at 1.4 bar (20 psi). Filters were placed into 50 µl conical tubes with 300 µl 1× PBS, 1.5 µl 2 % azide, and a sterile scalpel blade. Samples were minced using an OMNI bead ruptor elite (OMNI International, GA, USA) on a 4 m $s^{-1}$ 30 sec program. The resulting sample was centrifuged to collect bacteria separated from the filters and extracted for gDNA using a modified version of the Omega BioTek Universal Metagenomics kit protocol (OMEGA Bio-tek, GA, USA). Fifteen µl of MetaPolyzyme (Millipore Sigma, MA, USA) was added to each sample and incubated at 35 °C for 13 h followed by three cycles of freeze and thaw alternating between 80 °C for 2 min and a -80 °C freezer for 10 min. Further digestion was subsequently performed by adding 35 µl Proteinase K (Omega Bio-tek, GA, USA) and incubated at 55 °C for 1 hour. Following complete enzymatic digestion, the sample was extracted using the manufacturer's protocol (Omega BioTek Universal metagenomics kit). Briefly, 500 µl ML1 buffer (CTAB) was added the digested sample and incubated at 55 °C for 15 min. One volume of Tris-stabilized (pH >7.5) phenol-chloroform-isoamyl alcohol mix (25:24:1) was used for purification and the resulting upper aqueous phase was removed and combined with RBB (*Guanidinium) buffer and* 100 % EtOH and applied to a DNA silica column supplied with the kit. Final DNA was eluted in 35 µl of elution buffer. DNA was quantified using a Qubit spectrofluorometer and Nanodrop ND-1000 (ThermoFisher Waltham MA USA).

#### *RNA extraction*

For RNA extraction, total nucleic acids were extracted from polycarbonate filters (Millipore, 0.2 µm pore size) upon which 50 ml of ethanol fixed were collected. Extraction was done following Nercessian et *al.* (2005) with minor modifications. In brief, a CTAB extraction buffer containing SDS and N-laurylsarcosine was added to the samples together with an equal volume of phenol/chloroform/isoamylalcohol (25:24:1) solution. The samples were subject to bead-beating (FastPrep-24™ 5G Instrument, MP Biomedical, Eschwege, Germany), followed by centrifugation (14,000 *g*), a cleaning step with chloroform, and nucleic acid precipitation with PEG-6000 (Sigma-

Aldrich, Taufkirchen, Germany). The precipitated nucleic acids were rinsed with 1 mL of 70% ethanol, dried and dissolved in water. DNA was digested by two sequential treatments with the TurboDNAfree Kit (Invitrogen ThermoFisher Scientific, Dreieich, Germany) following the manufacturer's instructions. DNA removal was evaluated using a PCR reaction for 16S rRNA. First strand cDNA was then generated using the High-Capacity cDNA Reverse Transcription Kit (Applied Biosciences, ThermoFisher scientific), and was sent for sequencing at the Core Genomic Facility at RUSH university, Chicago, IL, USA.

#### *16S rRNA gene amplicon sequencing and processing*

PCR reactions were performed in triplicate. Each 25 µl reaction consisted of 0.5 µl of Phusion Green Hot Start II high-fidelity DNA polymerase (Thermo Fisher Scientific, Sweden), 5.0 µl of 5× Phusion Green HF buffer, 4 µl DNase- and RNase-free water, 5.0 µl of 10 µM primer mix (1:1), 0.5 µl of 10 mM nucleotide mix and 10 µl of DNA extract (0.5 ng/µl). The PCR protocol was 98 ºC for 30 sec, 33 cycles of 98 ºC for 10 sec, 55 ºC for 30 sec, 72 ºC for 30 sec and a final 5-min extension at 72 ºC. The amplicon target was the V3-V4 region of the 16S rRNA gene, using the V3 forward primer S-D-Bact-0341-b-S-17, 5'-CCTACGGGNGGCWGCAG-3 (Herlemann *et al.*, 2011), and the V4 reverse primer S-D-Bact-0785-a-A-21, 5'-GACTACHVGGGTATCTAATCC-3 (Muyzer *et al.*, 1993), resulting in fragments of ~430 bp. The primers were dual barcoded in a way compatible with Illumina sequencing platforms (as described in Caporaso *et al.*, (2011). Product size and successful amplification was tested by running internal positive and negative controls from the triplicate plates on a 1.5 % (w/v) agarose gel. Triplicate PCR products were combined, and each triplicate sample was purified using SPRI beads (Agencourt® AMPure® XP, Beckman Coulter, CA, USA). DNA concentration of purified samples was determined with a Quant-iT high-sensitivity DNA assay kit and a Qubit® fluorometer (Invitrogen, Carlsbad, USA). All samples were diluted to similar concentrations prior to pooling diluted PCR products together in equimolar volumes (50 µl) in one composite sample (including positive and negative controls).

Composite samples were paired-end sequenced at the Vrije Universiteit Amsterdam Medical Center (Amsterdam, The Netherlands) on an Illumina MiSeq Sequencer with a 600-cycle MiSeq Reagent Kit v3 (Illumina, San Diego, Ca, USA) according to the manufacturer instructions. Sequences were quality trimmed using Trimmomatic (v 0.39) and paired using Bbmerge (Bushnell *et al.*, 2017). The paired sequences were dereplicated using the dedupe tool of the BBTools package (sourceforge.net/projects/bbmap/) aligned and annotated using the SINA aligner (Pruesse *et al.*, 2012) against the SILVA SSU database (v 138.1) (Quast *et al.*, 2013)

A maximum-likelihood phylogenetic tree was calculated including only long 16S rRNA sequences, using FastTree 2 (Price *et al.*, 2010) using all *Thiovulum* sequences in the SILVA database (n=71), three 16S rRNA sequences obtained from the assembled genome of the Movile Cave *Thiovulum* (see below) and sequences of 4 different *Sulfurimonas* species as an outgroup. A second tree included amplicon sequences as well. For the sake of legibility, the 908 *Thiovulum* sequence variants obtained were clustered at 97 % similarity using CD-HIT-EST (Huang *et al.*, 2010), resulting in 50 clusters.

To obtain information on relative *Thiovulum* abundance, the raw short-read libraries (metagenomic) were analyzed with phyloFlash (V 3.3; (Gruber-Vodicka *et al.*, 2020)).

#### Shotgun sequencing (Illumina and Oxford Nanopore)

Shotgun sequencing was accomplished using both Illumina and Oxford Nanopore sequencing technologies. For Illumina sequencing, 1 ng of genomic DNA from each sample was converted to whole-genome sequencing libraries using the Nextera XT sequencing reagents according to the manufacturer's instructions (Illumina, San Deigo CA). Final libraries were checked for library insert size using the Agilent Bioanalyser 2100 (Agilent Technologies Santa Clara, CA) and quantified using Qubit spectrofluorometry. The final sample was sequenced using paired end 2x150 sequencing on an Illumina MiniSeq system.

A first pass of Oxford Nanopore sequences was obtained using the SQK LSK109 ligation library synthesis reagents on a Rev 9.4 nanopore flow cell with the GridION X5 MK1 sequencing platform, resulting in a total of 131.8 Mbp of reads with a N50 of 1.3 kbp.

Additionally, sequencing was performed on several cellular aggregates that were confirmed microscopically to contain *Thiovulum* cells. For this, the aggregates were picked from environmental samples fixed in self-made RNAlater (4 M $(NH_4)_2SO_4$; 15 mM EDTA (from 0.5 M, pH 8.0 stock); 18.75 mM Na-citrate (from 1M stock)), and washed several times in sterile 1× PBS buffer (137 mM NaCl; 2.7 mM KCl; 10 mM $Na_2HPO_4$; 1.8 mM $KH_2PO_4$; pH 7.4). The cell aggregates were lysed by freeze thawing and further, following the manufacturer's instructions, as part of the DNA amplification process using the Repli-G single cell amplification kit (Qiagene, Hilden, Germany). Libraries for Nanopore sequencing were prepared using the LSK-108 kit following the manufacturer's protocol but skipping the size selection step. The prepared libraries were loaded on MIN106 R9 flow cells, generating a total of 5.7 Gbp of reads with a length N50 of about 3.7 kbp. Basecalling for all Oxford Nanopore reads were done using Guppy 4.0.11.

#### cDNA sequencing

Provided cDNA was adjusted to 45 µl with water and sheared with the Rapid Shear gDNA shearing kit (Triangle Biotechnology, Durham, NC, USA). Briefly, 5 µ of Rapid Shear Reagent was added to the bottom of a 24-well PCR plate. cDNA (45 µl) was then added to the well and the plate was sealed and held on ice till shearing. Shearing took place for 5 minutes in a sonication bath. The resulting DNA at,of about 350bp isis length, was used directly in the Swift 1S protocol (Accel-NGS 1S Plus kit, Swift Biosciences, Ann Arbor, Mi, USA). The resulting sheared DNA was adjusted to 5ng/µ and a total of 75 ng was used for library prep, except for sample LR1 (Lake Room) that had low cDNA concentration. For this sample, the maximum volume of 15 µl was used as input. Library prep was as per the Swift Protocol with 6 cycles of PCR during indexing. Following library prep, all libraries were pooled in equal volume by combining 2 µl of each library for a final bead clean up with 0.85X AmpPure beads (Beckman Coulter Life Sciences, Indianapolis, IN, USA). This QC pool was then sequenced on an Illumina MiniSeq MO flow cell. The resulting index distribution was used to re-pool the libraries for an Illumina SP flow cell sequencing run with sample LR1 pooled at maximum volume available.

All sequencing data generated in this study were deposited in NCBI Sequence Read Archive under accession number PRJNA673084.

#### Metagenomic data analysis

Nanopore reads were assembled using Flye 2.8.1-b1676 (Kolmogorov *et al.*, 2019) with default parameters. The resulting GFA was examined using Bandage (Wick *et al.*, 2015), allowing to delineate a set of high-coverage (>300X) contigs against a background of low-coverage (<100X) contigs. To verify that these high-coverage contigs corresponded to *Thiovulum*, the published proteome of *Thiovulum* ES (Marshall *et al.*, 2012). was aligned on the GFA using tblastn (Gertz *et al.*, 2006) within Bandage (parameters: minimum identity 70 %, minimum coverage 70 %), revealing that nearly all tblastn hits were concentrated on the high-coverage contigs and vice-versa. The GFA was therefore pruned to retain only the high-coverage contigs, which were all interconnected. The remaining 31 contigs were exported as FASTA then scaffolded using SLR (Luo *et al.*, 2019); the nine resulting scaffolds were mapped back to the GFA to resolve most repeats, and the remaining repeats were resolved manually until obtaining a circular genome. A final polishing step was performed with unicycler-polish from Unicycler v0.4.9b (Wick *et al.*, 2017) using the complete set of Illumina reads (for a total depth of coverage of 12X of the genome) and the subset of Nanopore reads longer than 5 kb (*ca*. 50X). Polishing consisted of two cycles of pilon 1.23 (Walker *et al.*, 2014), one cycle of racon 0.5.0 (Vaser *et al.*, 2017) followed by FreeBase (Garrison and Marth, 2012), then 30 additional cycles of short-read polishing using pilon 1.23, after which the assembly reached its best ALE score (Clark *et al.*, 2013).

The completeness of the *Thiovulum* genome obtained was assessed using CheckM (Parks *et al.*, 2015) and its continuity using the unicycler-check module in Unicycler v0.4.9b. Annotation was performed using the command-line Prokka (Seemann, 2014) and DRAM (Shaffer *et al.*, 2020) tools as well asthe KEGG (Kanehisa *et al.*, 2016), EggNOG 5.0 (Huerta-Cepas *et al.*, 2019), PATRIC (Davis *et al.*, 2020;

Brettin *et al.*, 2015) and RAST (Aziz *et al.*, 2008; Overbeek *et al.*, 2014) annotation servers. A COG (Tatusov *et al.*, 2000) analysis was done using the ANVIO tool (Eren *et al.*, 2015). OperonMapper (Taboada *et al.*, 2018) was used to inspect the organization of genes into operons. CRISPRs where identified using CRISPR finder tool (Grissa *et al.*, 2007). To further elucidate the function of genes annotated as hypothetical proteins, a structural annotation was perforled using the Superfamily and SCOP databases using the Superfam online interface (Gough *et al.*, 2001). Metabolic models of the annotated genome from Movile Cave and that of *Thiovulum* ES were calculated using PathwayTools (V25.3)(Karp *et al.*, 2021).

#### *Thiovulum* sp. genome assembly from public databases

To obtain genomic information from additional cave-dwelling *Thiovulaceae* we downloaded all available metagenomic libraries from the Frasassi caves in Italy (SRR10997432, SRR1559028, SRR1559230, SRR1559353, SRR1560064, SRR1560266, SRR1560848, SRR1560849, SRR1560850, SRR8191123, SRR8194889, SRR8197024, SRR8200784, SRR8202337, SRR8203764). The short read libraries were quality-trimmed using Trimmomatic (Bolger *et al.*, 2014) and scanned for the presence of *Thiovulum* 16S rRNA using PhyloFlash (Gruber-Vodicka *et al.*, 2020), revealingthat library SRR1560850 contained >170,000 *Thiovulum* sp. 16S rRNA. A metagenomic assembly of library SRR1560850 wastherefore conducted using Megahit (Li *et al.*, 2015), after which the assembly was binned using Metabat2 (Kang *et al.*, 2015). The obtained bins were taxonomically annotated using the GTDB-Tk tool (Chaumeil *et al.*, 2019) resulting in 1Thiovulum1Thiovuluma single bin. The phylogenetic tree generated by the GTDB-Tk tool from a single-copy marker gene multilocus alignment suggested that the Movile and Frasassi caves *Thiovulum* genomes were closely related, hence, both genomes were used to recruit all *Thiovulum* related reads from all Frasassi libraries. The obtained reads were re-assembled, binned and taxonomically annotated, as above, resulting in two bins, one of which annotated as *Thiovulum* with 94 % completeness, 0.41 % contamination and 25 % strain heterogeneity as evaluated using CheckM (Parks *et al.*, 2015).

#### Transcriptomic analysis

The 6 libraries containing cDNA sequences (3 from Air Bell 2 and 3 from Lake Room), were quality-trimmed using trimommatic (Bolger *et al.*, 2014) and mapped against the complete genome of the Movile cave *Thiovulum* sp. using Salmon version 1.6 (Patro *et al.*, 2017). Ribosomal RNA data were removed from the mapping results and TPM (Transcripts Per Kilobase Million) values were recalculated to reflect mRNA expression. Further removal of tRNA sequences did not alter the results. The RNA data was analyzed using the iDEP (v. 0.95) online tool (Ge *et al.*, 2018) that provides an online graphical user interface for the DeSEQ2 (Love *et al.*, 2014) and Limma (Ritchie *et al.*, 2015) packages for RNAseq analysis. Differential expression was considered significant with a 2-fold difference and a false discovery rate smaller than 0.1. Taxonomic composition of the active community was obtain by analyzing the 16S rRNA gene from the transcriptomic read libraries using PhyloFlash as above (V 3.3; (Gruber-Vodicka *et al.*, 2020)). Viral transcripts were identified using VirSorter2 (v.1.1) (Guo *et al.*, 2021), annotated against the viral refseq database release 209 using BLAST and quantified using Salmon version 1.6 (Patro *et al.*, 2017).

Supplementary Figures

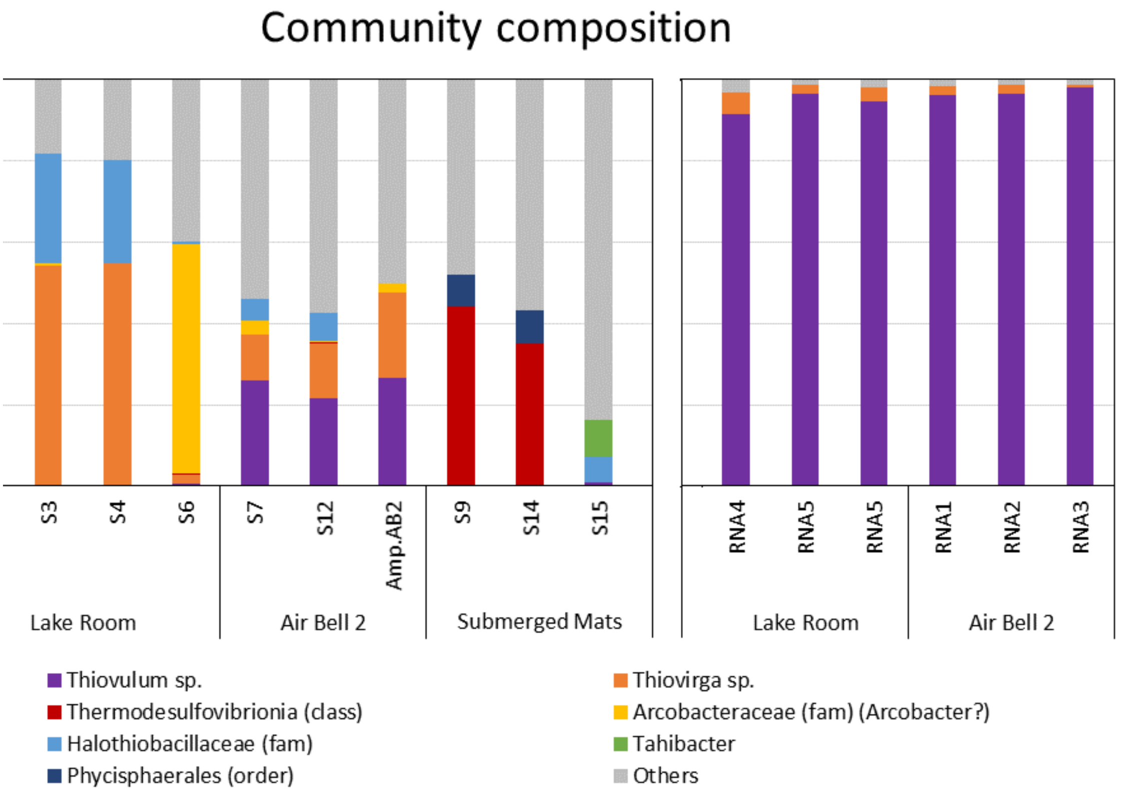


**Figure S1**. Community composition extracted from shotgun metagenomic reads (and one 16S rRNA amplicon sample) from Lake Room, Air Bell 2 and submerged microbial mats in Movile Cave. A long-term study on the cave microbial community based on amplicon sequences will be published separately.

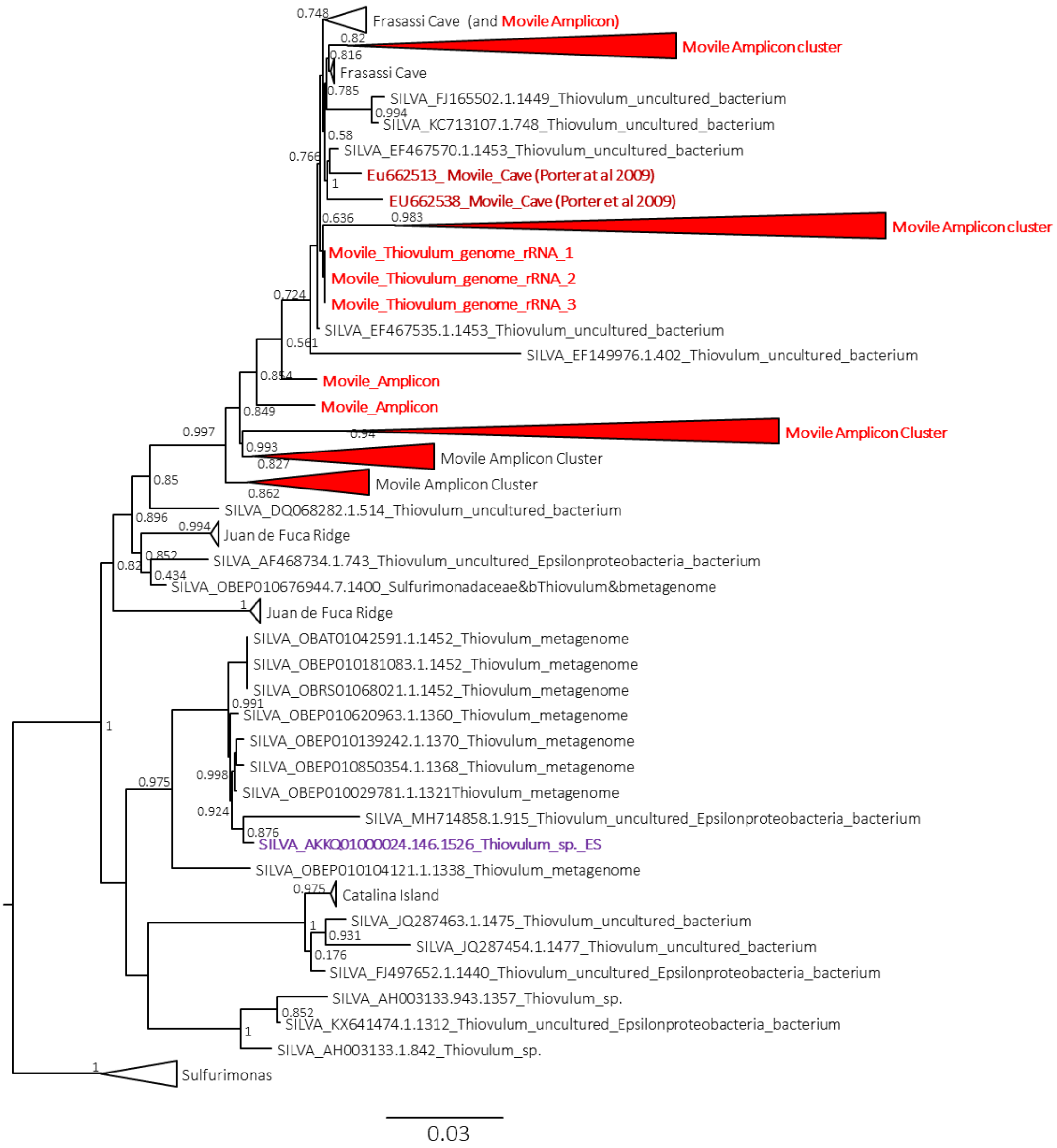


**Figure S2**. Maximum-likelihood placement of the 16S rRNA gene of the Movile Cave *Thiovulum* from Movile Cave. The 16S rRNA sequence in the genome of the Movile Cave *Thiovulum*, amplicons from Movile Cave, and all *Thiovulum* sequences available in the SILVA database (V138.1 Quast et al 2013) using *Sulfurimonas* sp. as an outgroup. The Shimodaira-Hasegawa local support values (ranging from 0 to1) are shown next to each node.

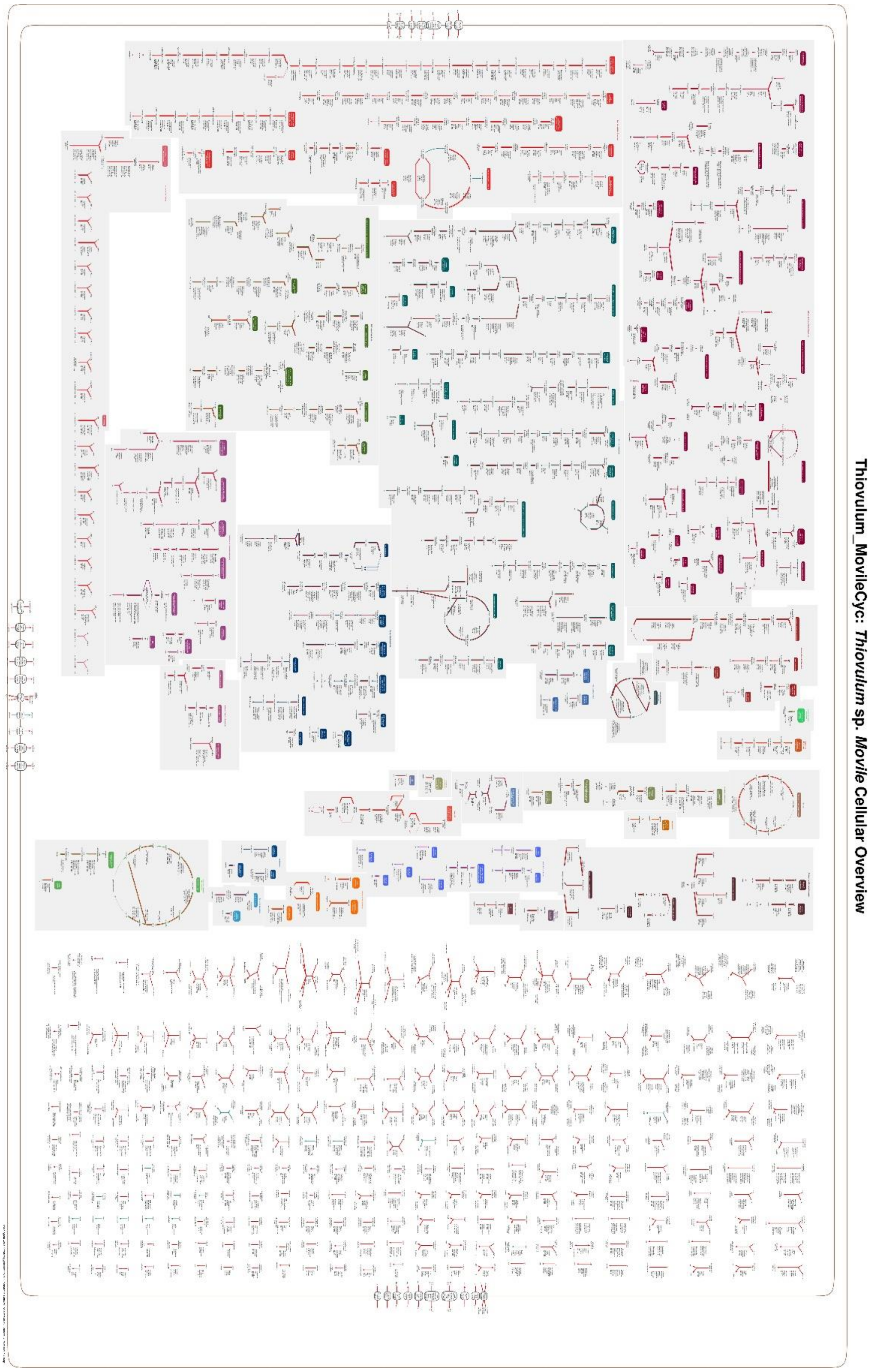


**Figure S3.** Overview of Movile *Thiovulum* metabolism Red lines mark commonalities with *Thiovulum* ES. Image will be separate and not part of a document.

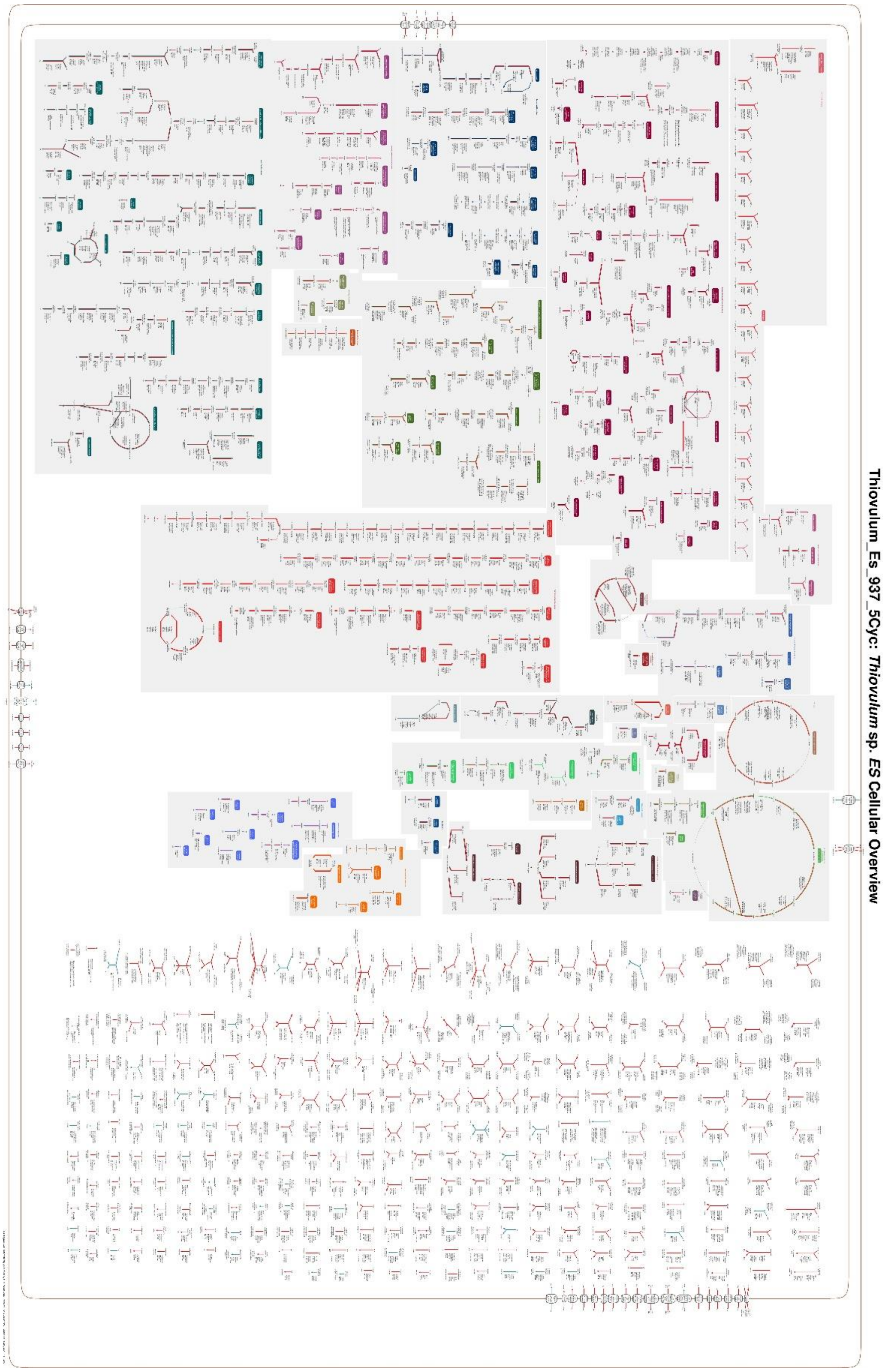


**Figure S4.** Overview of *Thiovulum* ES metabolism Red lines mark commonalities with Movile *Thiovulum*. Image will be separate and not part of a document.

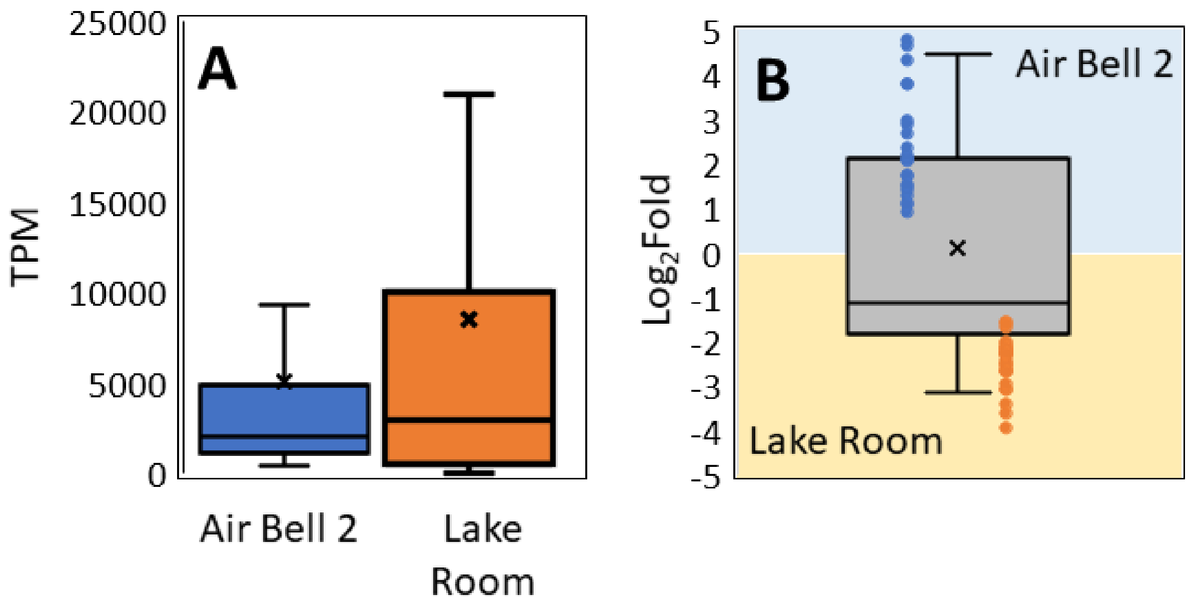


Figure S5. Quantification of reads from Air Bell 2 and the Lake Room mapping to viral transcripts expressed as transcripts per million (TPM) (A). Log 2 fold expression of viral transcripts in Air Bell 2 vs the Lake Room. Values above 1 and below -1 show a 2 fold change in expression in favor of Air Bell 2 and the Lake Room, respectively (B). The average line in panel B, close to a value of -1 highlights the higher expression of viral transcripts in the Lake Room.
